# Transcriptomic landscape of posterior regeneration in the annelid *Platynereis dumerilii*

**DOI:** 10.1101/2023.05.26.542455

**Authors:** Louis Paré, Loïc Bideau, Loeiza Baduel, Caroline Dalle, Médine Benchouaia, Stephan Q. Schneider, Lucie Laplane, Yves Clément, Michel Vervoort, Eve Gazave

## Abstract

**Background:** Restorative regeneration, the capacity to reform a lost body part following amputation or injury, is an important and still poorly understood process in animals. Annelids, or segmented worms, show amazing regenerative capabilities, and as such are a crucial group to investigate. Elucidating the molecular mechanisms that underpin regeneration in this major group remains a key goal. Among annelids, the nereididae *Platynereis dumerilii* (re)emerged recently as a front-line regeneration model. Following amputation of its posterior part, *Platynereis* worms can regenerate both differentiated tissues of their terminal part as well as a growth zone that contains putative stem cells. While this regeneration process follows specific and reproducible stages that have been well characterized, the transcriptomic landscape of these stages remains to be uncovered.

**Results:** We generated a high quality *de novo* Reference transcriptome for the annelid *Platynereis dumerilii*. To do so, we produced and analyzed three RNA-sequencing datasets, encompassing five stages of posterior regeneration, along with blastema stages and non-amputated tissues as controls. We included these regeneration RNA-seq datasets, as well as embryonic and tissue-specific datasets from the literature to produce a Reference transcriptome. We used this Reference transcriptome to perform in depth analyzes of RNA-seq data during the course of regeneration to reveal the important dynamics of the gene expression, process with thousands of genes differentially expressed between stages, as well as unique and specific genes expression at each regeneration stage. The study of these genes highlighted the importance of the nervous system at both early and late stages of regeneration, as well as the enrichment of RNA-binding proteins (RBPs) during almost the entire regeneration process.

**Conclusions:** In this study, we provided a high-quality *de novo* Reference transcriptome for the annelid *Platynereis* that is useful for investigating various developmental processes, including regeneration. Our extensive stage-specific transcriptional analysis during the course of posterior regeneration shed light upon major molecular mechanisms and pathways, and will foster many specific studies in the future.

## Background

Restorative regeneration, the ability to reform a lost or damaged body part, is a fascinating morphogenetic process that has intrigued scientists for centuries (1, 2). Among Metazoa, this capability is shared by many animal lineages, from non-bilaterians (sponges, cnidarians, ctenophores and placozoans) to mammals (including humans), but it varies greatly in scales. Indeed, while the majority of animal phyla possesses at least one species that is able to regenerate an organ (with the exception of few lineages of Ecdysozoa), fewer are those that can regenerate a complex structure, such as a limb (*i.e.* salamander) or a body axis (*i.e.* annelid) (2, 3). The extreme and ultimate regeneration ability, whole-body regeneration from a small piece of tissue, is even more restricted and remains the prerogative of non-vertebrates (non-bilaterians, some spiralians, as well as few deuterostome groups) (2, 3). In addition, regeneration abilities also vary from one tissue to another and during the life cycle of a given species (2, 4). Notwithstanding this huge diversity in regeneration modalities, it is worth to note that almost all regeneration processes can be virtually divided in three sequential steps shared by all regeneration model systems. The first one, wound healing, corresponds to the wound closure through the reconstitution of an epithelium. The second step usually relies on the formation of a regeneration-specific structure, the blastema, a result of the mobilization of precursor cells. The last step encompasses the morphogenetic processes that involves patterning, differentiation and growth of the reformed structure (2, 5, 6).

Despite its crucial role in animal life cycles, the fundamental process of regeneration remains poorly understood, and elucidating its molecular mechanisms is still a lively question (7, 8). One way to approach is to identify genes that are activated during the main steps of this process. Transcriptomic data, notably staged-bulk RNA-seq datasets, have shown to be an efficient way to achieve that. Major advances in the regeneration field were accomplished due to transcriptomic studies. These include studied targeting long-lasting questions regarding i) the extant of recapitulation of development during regeneration and its fidelity (9–11), ii) the (re)patterning and integration of a newly (re)formed structure (12, 13), iii) the variation of regenerative capabilities (14, 15) and iv) the evolutionary history of regeneration (2, 16).

Among animal lineages showing extensive regenerative capabilities, annelids, or segmented worms, are a historically important phylum to be investigated (17–22). Annelids comprise a large group of around 22 000 marine, freshwater and terrestrial species (23, 24) mainly composed of two major subdivisions, the Errantia and the Sedentaria (25, 26). Most annelids, from both subdivisions, harbors important regenerative capabilities, with the exception of leaches (Sedentaria, Clitellata) that are almost devoid of any regenerative ability (17, 18, 27). Following amputation or injury, many annelids are able to reform the posterior part of their body and, for many families, their anterior one including the entire head. Annelids are also able to regenerate various appendages, notably their crawling ones named parapodia, as well tentacles and cirri associated with terminal regions or body segments. While the ability to regenerate the posterior parts and appendages is shared between all regeneration-competent annelids, anterior or head regeneration is more restricted (17, 28). Currently, several species (such as *Pristina leydyi, Capitella teleta, Enchytraeus japonensis* …), all members of the Sedentaria clade, are studied to answer specific questions about annelid regeneration, for example, the different sources of cells involved in the blastema formation (29–33). In addition, due to technological advances in next generation sequencing, we slowly start to uncover transcriptional profiles of annelid regeneration. These include Bulk RNA-seq data from several Clitellata species during either posterior or anterior regeneration (34–36). A recent study compared transcriptional profiles of anterior and posterior regenerations with posterior growth in syllids (Errantia), suggesting that posterior growth and regeneration are more similar to each other than the two regeneration processes are (37). Despite these progresses, a broad basic understanding and description of the regenerative processes is lacking for those species and few molecular and functional tools, - if any -, are available. In conclusion, while annelids show important regenerative capabilities and have a rich history of experimental studies, we are still lacking a front-line regeneration model species to address fundamental and mechanistic questions about regeneration, amenable for molecular and functional studies.

One key annelid model species, developed for decades with success, is the Nereididae (Errantia) *Platynereis dumerilii* (38). *Platynereis* has risen as an intensely studied model organism for developmental, marine, neural, and evolutionary biology, in which a variety of technical tools are available (24, 39). *Platynereis* (re)emerged as an interesting regeneration model system over the past few years as it shows extensive regenerative capabilities (28). After amputation of the posterior part of their body, which leads to the removal of the pygidium (terminal non-segmented body part of the worm), the stem cell-rich sub-terminal growth zone (responsible for the continuous growth of the worms (40)) and several segments, *Platynereis* worms are able to quickly and properly regenerate their posterior part quickly (41) (Additional file 1). Briefly, after amputation (stage 0 or 0-day post-amputation (dpa)), a wound-healing epithelium closes the wound within 24 hours (stage 1 or 1 dpa). One day later (stage 2), the blastema starts to form. Those two steps do not involve much cell proliferation, which is non-mandatory. At 3 dpa (stage 3), the blastema enlarges and extensive cell proliferation is observed. Between stages 2 and 3, the majority of the tissues and structures (*i.e.* muscles, segments, nervous system …) begin to reestablish. At stage 5 (5 dpa), the growth zone is functional and have started to produce a few segments, while still not morphologically visible yet. As such, at this stage, the regeneration process is finished and posterior growth has resumed (41) (Additional file 1). In terms of genomic and transcriptomic resources: there is currently no published *Platynereis dumerilii* genome, while some transcriptomic data has been essentially generated from embryonic and larval stages (42–44), as well as from brain tissue (45). Few scRNA-seq datasets from early embryos and larvae have also been generated (46, 47). A major gap remains in the exploration of the molecular machinery involved in post-embryonic processes, especially regeneration.

In this study, we generate a new high-quality transcriptome of *Platynereis dumerilii*, encompassing both embryonic, larval and regeneration data, that will be critical in the future for all kind of transcriptomic and functional studies in this model species. Here we utilize this new resource to characterize the transcriptomic landscape of posterior regeneration and highlight key molecular mechanisms that underlie this intricate process, focusing notably on transcription factors and RNA-binding proteins, signaling pathways as well as nervous system and stem cells gene markers.

## Results and discussion

### A high-quality Reference transcriptome suited for studying both regeneration and development in *Platynereis dumerilii*

#### Step-wise reconstruction of a reference transcriptome in P. dumerilii

The main aim of our study was to investigate the transcriptional landscape of posterior regeneration in an emerging regeneration model: the annelid *Platynereis dumerilii* (39). Hence, it was mandatory to reconstruct a high-quality reference transcriptome suitable for the study of this post-embryonic developmental process, as so far available transcriptomes are mostly encompassing early developmental stages or specific tissues (43–45). To this end, we took advantage of a variety of *Platynereis*’ data sources collected from the literature (44, 45), and specifically generated additional missing transcriptomic data.

We generated both short and long read RNA-seq data during the posterior regeneration process. Short reads (150bp, paired-end) were generated from two regeneration stages, 2 days post-amputation (dpa) and 3dpa, which correspond to the blastema formation ((41), Additional file 1), and will be referred to “Blastema” data (or series 1) for the rest of the manuscript. Long reads were generated from the blastema of regenerating posterior parts at 3dpa, as well as from non-amputated posterior parts, and will be referred to “long reads” data (or series 2) for the remainder of the manuscript (Additional file 2).

To reconstruct an optimal reference transcriptome in *P. dumerilii*, we decided to first reconstruct three intermediate transcriptomes, and second merge them along with two other transcriptomes from the literature (Additional file 3). We used two widely-used tools (namely Trinity and rnaSPAdes) as they are considered to perform best amongst transcriptome reconstruction tools (48). This experimental strategy has the advantage of combining the strengths of both tools and fully exploit all our sequencing datasets (short and long reads) in depth. Moreover, merging transcriptomes from different assemblies and sources using EvidentialGene gives rise to better quality assemblies (*e.g.* measured in the number of full-length protein-coding transcripts) and single-tool transcriptome assemblies, as noted in plants and mammals (49, 50).

We assembled a first transcriptome only using the series 1 using Trinity [50], referred to as the Blastema transcriptome. We assembled a second transcriptome using the series 1 (short reads) and long reads datasets (series 2) using rnaSPAdes (48), referred to as the Hybrid 1 transcriptome. Unlike short reads, long reads can easily span several exons, as a result, their integration in transcriptome assembly greatly improves assembly quality, notably in the context of alternative splicing. We assembled a third transcriptome using again the series 1 and 2 datasets, this time using Trinity, referred to as the Hybrid 2 transcriptome. We finally merged these three transcriptomes, along with two additional transcriptomes available from the literature. The first one, generated with Trinity from short read RNA-seq during development and encompassing seven early embryonic stages (from 2 to 14 hours post-fertilization), is referred to as the Embryonic transcriptome (44). The other one, also generated with Trinity, includes short read RNA-seq from the heads of male and female worms at different steps of sexual maturation, and is referred to as the Head transcriptome (45). We then generated a Reference transcriptome from the merging of these five transcriptomes (See Additional files 2 and 3).

#### Quality assessment of transcriptome assemblies

In order to evaluate the quality of the generated transcriptome, in comparison to the other ones, we computed classical metrics for all of them. The “Blastema”, “Hybrid 1”, “Hybrid 2”, “Embryonic”, “Head” and “Reference” transcriptomes contain respectively 382 090, 315 141, 536 225, 273 087, 52 059, and 331 646 transcripts (Additional file 4). These differences likely reflect the differences in data type and quality, as well as the reconstruction methods. Indeed, Trinity software is known to generate an artificially high number of transcripts (51). Regarding the “head” transcriptome which harbor a very low number of transcripts, this is due to an extremely stringent strategy implemented by the authors to eliminate redundancy (*e.g.* only keep the longest transcript in case of multiple copies, (45)). Although the number of transcripts in our Reference transcriptome is intermediate, it has the highest mean and median transcript lengths (1 890bp and 1 256bp respectively), the highest number of transcripts longer than 1 000bp, and the second highest number of transcripts longer than 10 000bp (Figure 1A, Additional file 4), which indicates gain of assembly quality with this Reference transcriptome as compared to formerly available transcriptomes. This also highlights that, as expected, the inclusion of long read data allowed to obtain of very large – potentially complete - transcripts, in comparison to Illumina reads (even 150 bp). Regarding the assemblers we tested, our analyses confirmed that rnaSPAdes is much more powerful than Trinity to integrate data of different types, *i.e.* Illumina short read and ONT (Oxford Nanopore Technologies) long-read, as expected based on previous studies (48).

**Figure 1:**
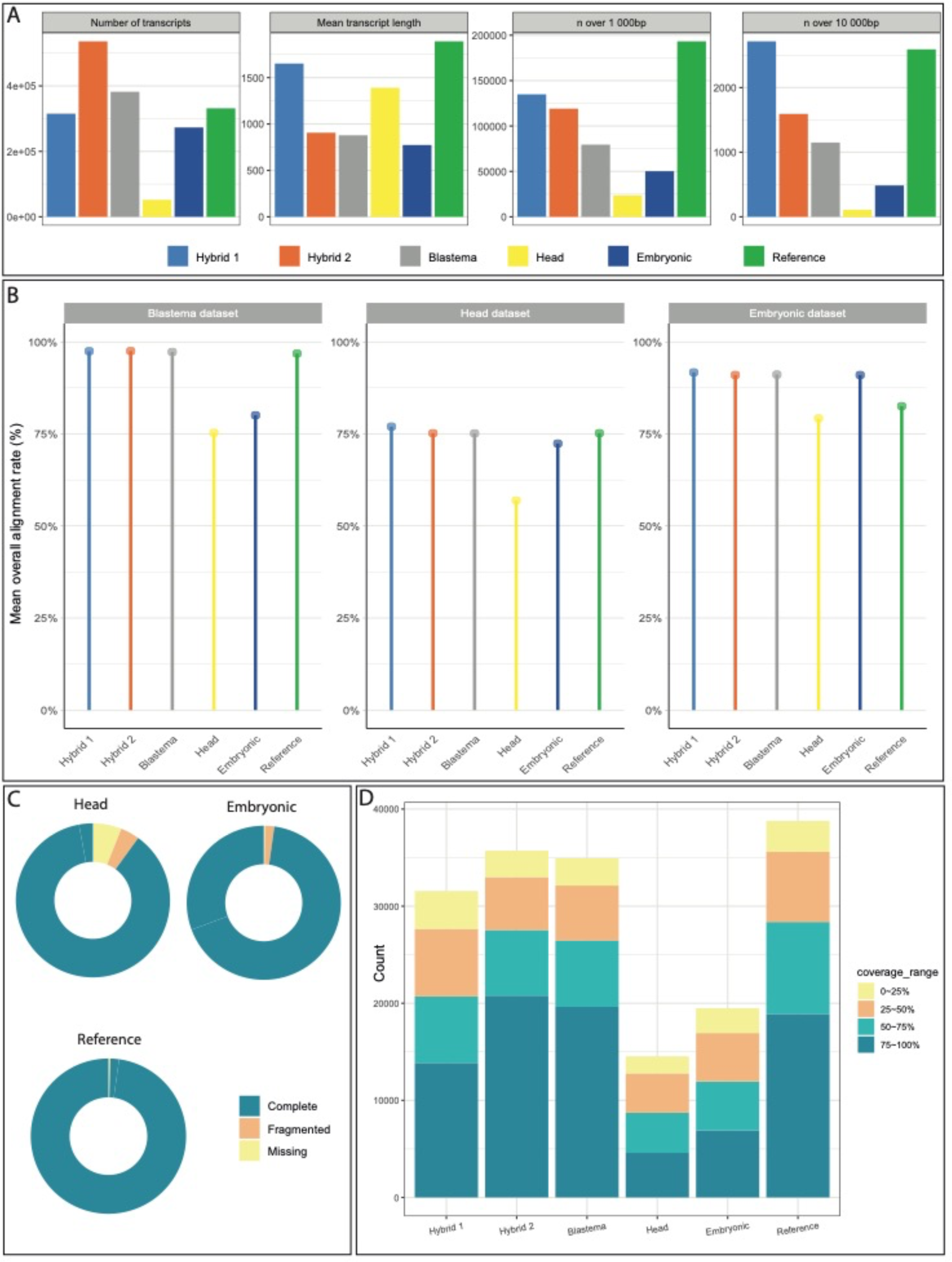
Quality control metrics for *Platynereis dumerilii* transcriptomes. A) Total number of transcripts (left), mean transcript length (middle left), number of transcripts with length above 1000bp (middle right) and above 10 000 bp (right) are shown. All transcriptomes are represented, including our intermediate assemblies (Hybrid 1, Hybrid 2 and Blastema), the two from the literature (Head and Embryonic), and our Reference assembly. B) Mean alignment rate (percentage of reads mapped) for each transcriptome assembly. Reads from the Blastema, the Embryonic and the Head datasets (see Additional file 2) were mapped on the six transcriptomes with bowtie2 (93). C) BUSCO completeness metrics for the Embryonic, the Head, and our Reference transcriptomes. Complete single-copy and complete duplicated were merged because of the redundancy that is normally occurring in de novo transcriptome assemblies. D) Swiss-Prot BLASTX results for each transcriptome assembly. Total bar height represents total number of transcripts with a significant BLASTX hit (e-value cutoff 10^-5^), colored bars represent the number of transcripts having the indicated coverage with a Swiss-Prot sequence. All BLASTX were done on the non-redundant Swiss-Prot database.

Moreover, we looked at reconstruction metrics, namely N50 (the length of the shortest transcript for which longer and equal size transcripts cover at least 50% of the transcriptome sequences) (52), and Ex90N50 (N50 for the 90% most expressed transcripts) (Additional file 4). N50 for the “Blastema”, “Hybrid 1”, “Hybrid 2”, “Embryonic”, “Head”, and “Reference” are 1730, 3565, 1869, 1466, 1822, and 2933, respectively. The N50 for the Reference transcriptome is not the best, but it’s the second highest after “Hybrid1”. Ex90N50 values, however, are highest for the Reference transcriptome (3093 compared to 3077 for the second highest, Additional file 4).

In addition, we wanted to assess how well various types of RNA-seq reads could align to the different transcriptomes we compared (Figure 1B, Additional file 5). The objective here is to have a Reference transcriptome that can be used in different biological contexts, from embryonic development to regeneration. Hence, we mapped RNA-seq reads from the “Blastema”, “Head”, and “Embryonic” datasets (Additional file 2) onto the six transcriptomes. Globally, the “Head” and “Embryonic” transcriptomes generally have the lowest mean percentage of mapped reads amongst the RNA-seq datasets that we investigated: 75% and 80% compared to at least 91% for other transcriptomes in the Blastema series, 57% and 72% compared to at least 75% for the Head series (Figure 1B, Additional file 5). The Embryonic transcriptome, however, has one of the highest mapping percentages for the Embryonic series (91%, (Figure 1B, Additional file 5), which can be explained by the fact that the Embryonic transcriptome was reconstructed from the RNA-seq reads of the Embryonic series. Surprisingly, the Head transcriptome exhibits the lowest mapping percentage even for reads from the Head series (57%, (Figure 1B, Additional file 5), despite the fact that this transcriptome was reconstructed using these reads specifically. We think this is due to the extreme low number of transcripts present in this assembly, because of stringent filters and parameters used by the authors when reconstructing the transcriptome. We note that our “intermediate” assemblies (namely “Hybrid 1”, “Hybrid 2” and “Blastema”) have the highest mapping percentages amongst all assemblies (Figure 1B, Additional file 5), while the Reference transcriptome has intermediate mapping percentages, between “intermediate” assemblies and “external” assemblies (namely “Head” and “Embryonic”). This was expected, as the Reference transcriptome is a mix between our “intermediate” assemblies and the “external” assemblies (Figure 1B, Additional file 5).

Taken together, these results indicate that, non-unexpectedly, transcriptomes assembled from reads having a single biological origin (*e.g.* “Blastema” or “Embryonic”) achieve highest mapping percentage with reads from the same biological origin (with the notable exception of the “Head” transcriptome, as mentioned before). This will cause some issues when using a reference transcriptome corresponding to a particular biological origin to map sequencing reads from a different origin (*e.g.* reads generated when studying regeneration mapped on an “embryonic” transcriptome). It is thus necessary to have a transcriptome assembled from the largest possible number of sources, thus reflecting the various biological processes that researchers are studying. This is what we’re achieving with our Reference transcriptome. As such, even though it harbors lower mapping percentages than the tissue/stage-specific assemblies, we argue that our Reference transcriptome represents the right balance between biological information and mapability.

We then determined the completeness of the transcriptomes using BUSCO. Using either the Eukaryota, Metazoa or Arthropoda conserved single-copy orthologs gene sets, the Reference transcriptome has the highest BUSCO scores (from 98.2 to 100 %) compared to other assemblies (Figure 1C and Additional file 6). We also looked at how well assemblies could recapitulate known *P. dumerilii* transcripts that are present on the NCBI database at the time of our analyses (n=13554). The Reference transcriptome has the highest recall rate for known transcripts, with more than 99% of *P. dumerilii* transcripts having a significant BLAST hit, compared to 93% for the Head and almost 98% for the Embryonic transcriptome (Additional file 6). This clearly indicates the high consistency between our Reference transcriptome assembly and the already known *Platynereis* genes.

We finally analyzed how well our transcripts recapitulated known proteins using BLAST analyses on Swiss-Prot database (Figure 1C and Additional file 6). Our Reference transcriptome has the highest percentage of transcript with at least one significant hit (e-value cutoff of 10^-5^) with 54% of transcripts, against 14% and 37% of transcripts for the Embryonic and Head transcriptome, respectively, and 32%, 33% and 23% for the Blastema, Hybrid 1 and Hybrid 2 transcriptomes, respectively. This result clearly indicates that merging transcripts from various biological conditions is generating a higher-quality transcriptome containing a more relevant biological information. Moreover, we looked at the coverage of query sequences (*i.e.* the percentage of *P. dumerilii* transcripts covered by a Swiss-Prot sequence, Fig. 1C and Additional file 6). Numbers, for a >75% coverage, indicate that this coverage is high but not the best for the Reference transcriptome (48 % *versus* 56 and 58 % for Hybrid 2 and Blastema transcriptomes, respectively; but also 44 % for Hybrid 1, 32 % for “Head” and 35.5 % for “Embryonic”). This is expected as transcripts in our Reference transcriptome are on average longer than in other transcriptomes, and consequently having a higher percentage of coverage is more complicated than for shorter transcripts (Fig. 1A, Additional file 4).

#### Functional annotation of the Reference transcriptome

We finally annotated the Reference transcriptome using both Trinotate (Additional files 7 and 8), or individual BLASTX searches on six specific species: four deuterostomes (*Homo sapiens*, *Mus musculus*, *Danio rerio*, and *Saccoglossus kowalevski*) and two protostomes (*Drosophila melanogaster*, and *Aplysia californica*) (Additional files 8 and 9). Trinotate results indicate that at least 57% of *P. dumerilii* transcripts have a significant BLASTX hit. Furthermore, we could find Pfam domains on 54% of all our transcripts and assign more than 50% of our transcripts to a KEGG pathway. While the percentage of KEGG-annotated transcripts is higher than reported in the “Embryonic” transcriptome, the percentage of transcripts with at least one Pfam domain is lower. However, this annotation in the “Embryonic” transcriptome was performed only on a subset of transcripts, namely those for which a predicted ORF longer than 100aa could be predicted (44), a restriction that we did not make for our Reference transcriptome.

BLAST searches in individual species show that between 38% and 53% of *P. dumerilii* transcripts have a hit in one of the six organisms, ranging from 38% in *D. melanogaster* to 53% in *D. rerio* (Additional file 9). These numbers are lower than what could be expected for an annelid species. Indeed, we performed the same BLAST analysis using transcripts of *Owenia fusiformis* a species with a well-sequenced and annotated genome (53) on the same species. This yielded higher figures, ranging from 59% for *D. melanogaster* to 73% for *S. kowalevskii*. Moreover, while around 80% of *O. fusiformis* transcripts had a significant hit (e-value cutoff value of 10^-3^) in a non-redundant database of annelid genes (53), 68% of *P. dumerilii* from our Reference transcriptome had such a significant hit. Although these numbers indicate that our Reference transcriptome assembly does not attain the quality of a reference genome, the (relatively) high figures we obtain with these BLAST analyses should be taken as an indication of the quality of our transcriptome assembly.

Taken together, these results show that our *de novo* Reference transcriptome is a high-quality transcriptome that probably represents the majority of genes that are transcribed during developmental events, with a high fraction of complete gene models and is thus well suited for the study of various biological processes of *P. dumerilii*.

### Transcriptomic landscape of *Platynereis’* posterior regeneration

Utilizing this new high-quality Reference transcriptome, we now have the opportunity to properly explore the transcriptomic landscape during posterior regeneration in *Platynereis dumerilii*. We previously defined key regeneration stages, at the morphological and cellular levels (41). For this bulk RNA-seq experiment, we selected five stages that encompass the entire regeneration process: stage 1 (1 dpa or wound healing step), stages 2 and 3 (2 and 3 dpa or blastema formation step) and stage 5 (5 dpa or last step of morphogenesis). For these stages, we collected the regenerated part, if any (*e.g.* there is only a regenerated epithelium at stage 1), as well as five upstream segments. A non-amputated stage, corresponding to the terminal part of the worm (*i.e.* the pygidium, the growth zone containing stem cells (40)) and the last five segments was also included. As a control, we generated samples at stage 0, (*i.e.* immediately after amputation or 0 dpa), collecting the five upstream segments (Additional file 1).

#### • Mapping, samples reproducibility and gene expression levels

We mapped this newly produced regeneration RNA-seq data (n=18 samples) to the Reference transcriptome we assembled. As for the other datasets, the percentage of successful mapping is very high (above 90% on average) ensuring that we can explore the huge majority of the generated data (Additional file 5).

We assessed the quality and reproducibility between our biological replicates through a Principal Component Analysis (PCA) and eliminated three samples that were identified as outliers (namely 1dpa_3, 3dpa_1 and 5dpa_3, Additional file 10). Consequently, for the 1, 3 and 5 dpa stages, our analyses only include two biological replicates, while we have 3 replicates for the NA, 0 and 2 dpa stages. In contrast to *Platynereis*’ embryonic developmental stages, which are extremely stereotypical (54), the regenerative stages, while well-defined (41), are a little less stereotypic and show a greater degree of biological variability. This may explain the presence of these outliers in our RNA-seq data. The PCA analysis also highlights the fact that 0, 1, 5 dpa and NA samples are relatively close together, while 2 and 3 dpa samples are very different, both from each other, and from other stages. We calculated the gene expression abundances for the high-quality samples (n=15).

We then evaluated the percentage of “genes” that was expressed in each of our conditions (out of a total of 119 737 genes). For that purpose, we selected three thresholds for TMM values. As for *Platynereis* embryonic data (44), we considered genes having a TMM > 1 as expressed. Because we also wanted to find how many genes were moderately or highly expressed, we defined two other thresholds at 5 TMM and 10 TMM. We considered genes with a TMM below 1 as not expressed, between 1 and 5 as being expressed, between 5 and 10 as moderately expressed and above 10 as highly expressed (Figure 2A, Additional file 12). Globally, all samples show similar proportions of genes in each category, with an average of 72% of genes that are not considered as expressed (0 < TMM <1). This high number of non-expressed genes is relatively expected for a transcriptome *de novo*, with thus an overestimated number of genes or transcripts. Around 17 % of genes are lowly expressed (1< TMM <5), 5% are moderately expressed and 5% are considered as highly expressed (Figure 2A).

**Figure 2:**
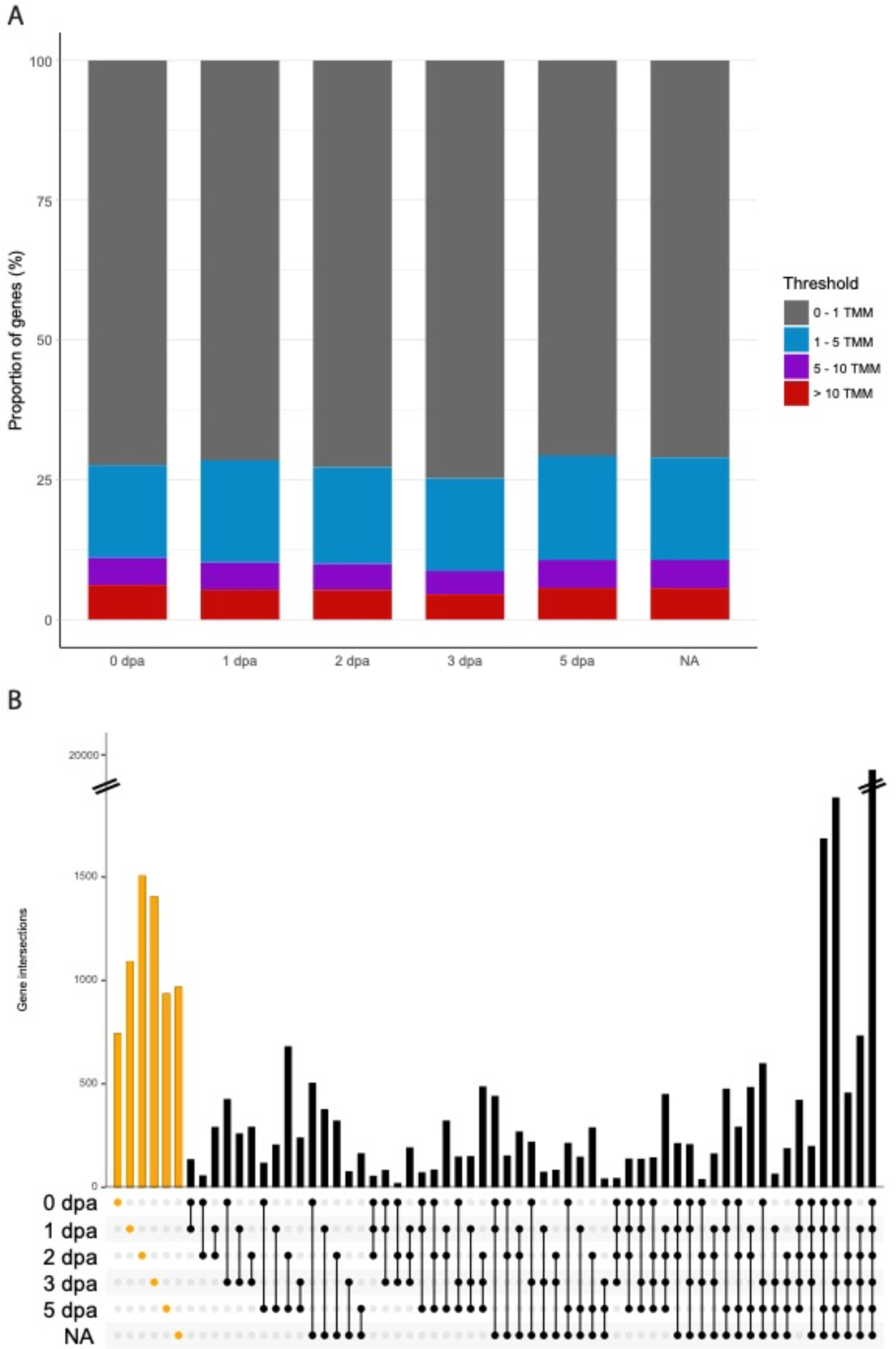
Global gene expression profiles during *Platynereis dumerilii* regeneration. A) Proportion of P. dumerilii transcripts with TMM values between 0 and 1 (grey), between 1 and 5 (blue), between 5 and 10 (purple), and above 10 (red) for five regeneration stages (0 dpa, 1 dpa, 2 dpa, 3 dpa, 5 dpa) and non-amputated control worms (NA). B) UpSetR plot (101) representing transcripts co-expressed (i.e. TMM value > 1) in various stage conditions. Bar heights represent the number of transcripts either expressed in only one condition (orange ones) or co-expressed in a number of conditions (black). All TMM values were computed from raw read counts using edgeR (99).

Then, we evaluated the number of genes that are expressed (TMM > 1) at a specific stage or across several stages by computing an upset plot (Figure 2B). Three key findings can be deduced from these results. First, the majority of genes (n=19 460 or 42 % of the 46039 unique genes with TMM>1) are expressed at every stage. Second, with the exception of these genes and two other combinations, genes expressed at only one stage represent the other largest categories (n=745, 1093, 1507, 1407, 937 and 969 for 0, 1, 2, 5dpa and NA, respectively). Third, genes expressed in a combination of stages only represent a small fraction of all *Platynereis* genes (Figure 2B).

#### • Global gene expression dynamics during posterior regeneration

We then aimed to dissect the dynamics of the transcriptional landscape during the course of regeneration. To this end, we first determined the genes that are differentially expressed (up or down regulated) between all pairs of regeneration stages (Additional file 13; Figure 3A), and could identify several thousands of such genes (Fig. 3A). We decided to focus our interest mostly on sequential conditions, as they have the most biological significance (*e.g.* 0 to 1 dpa, 1 to 2 dpa …). Regarding the genes that are upregulated from one stage to the next, their numbers vary from a minimum of 2619 (5 dpa *vs* NA) to a maximum of 8081 (3 *vs* 5 dpa). 4038 genes are significantly up-regulated between the amputation stage (0 dpa) and the wound healing stage (1 dpa).

**Figure 3:**
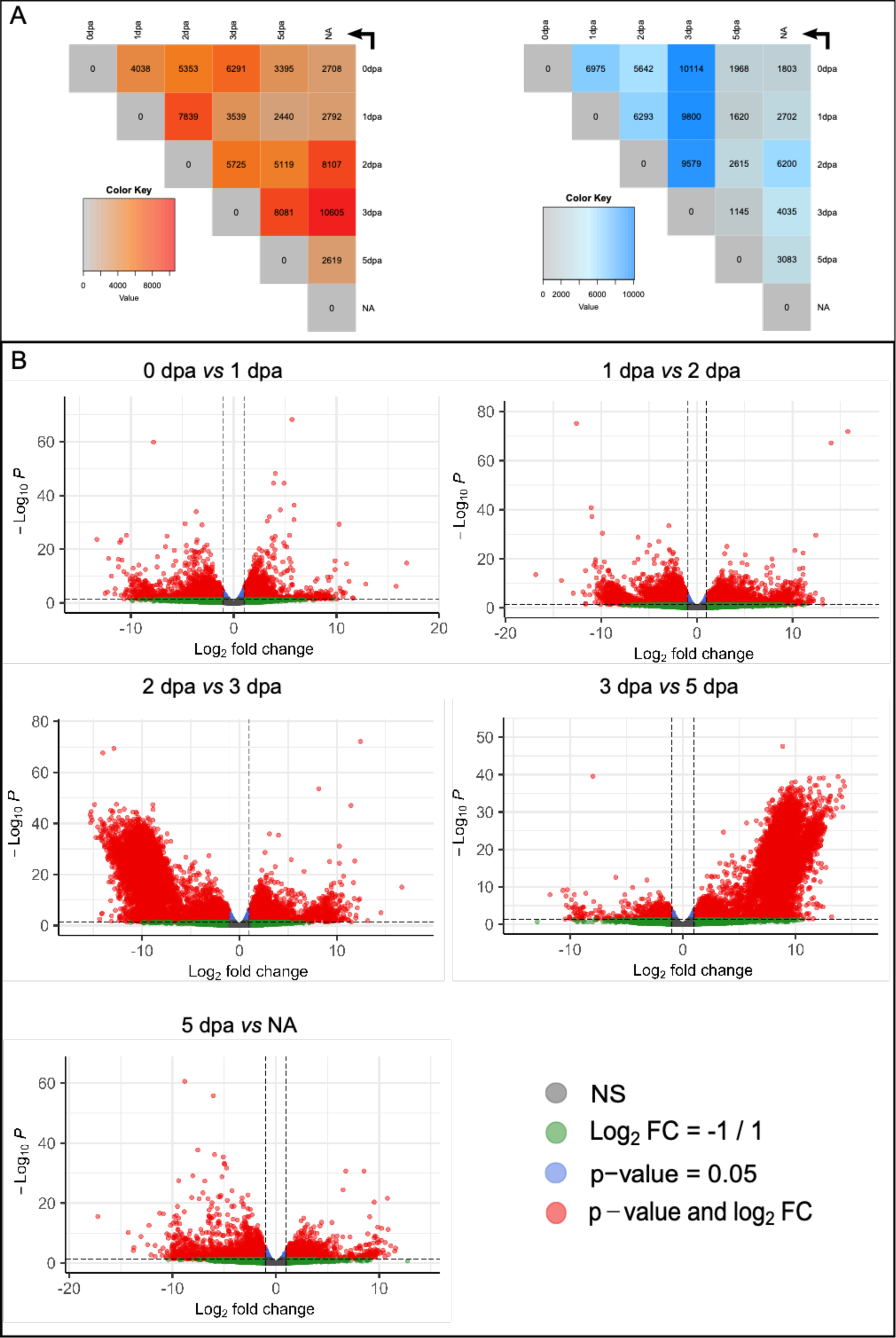
Differences of gene expression during regeneration in *Platynereis dumerilii*. A) Heatmaps showing differentially expressed genes (DEGs) between each pair of conditions, with up-regulated genes on the left (shades of red) and down-regulated genes on the right (shades of blue). Each line represents the control (or reference) conditions, each column represents the test conditions. For example, 3395 transcripts are up-regulated at 5 dpa compared to 0 dpa (as indicated by the arrow). The color key corresponds to the number of transcripts in each box. A transcript was considered as up-regulated (or down-regulated) with an FDR < 0.05 and a logFC > 1 (resp. < 1). DEGs were identified using edgeR (99). B) Volcano plots for sequential comparisons (0 dpa vs 1 dpa, 1 dpa vs 2 dpa, 2 dpa vs 3 dpa, 3 dpa vs 5 dpa, 5 dpa vs NA). Each dot represents a transcript, with colors indicating different values for FDR and logFC. In grey: FDR > 0.05; -1 < logFC < 1. In green: FDR > 0.05; logFC < -1 or logFC > 1. In blue: FDR < 0.05; -1 < logFC < 1. And finally, in red: FDR <0.05, -1 < logFC < 1. Thus, the DEGs are depicted in red.

These DEGs are potentially involved in the epithelium reformation step, as well as potential events taking place in segments abutting the amputation plane. Between 1 dpa and the blastema’s initiation stage (2 dpa), a very high number of DEGs is identified (7839), and may correspond to genes activated and required for blastema initiation. Between the two blastema stages (2 and 3 dpa), the number of up-regulated genes is 5725, revealing that during blastema growth, an important transcriptomic activity is still mandatory. We also observed a huge number of up-regulated genes (8081) between 3 and 5 dpa highlighting the fact that a blastema corresponds to a structure still in the course of differentiation, very different from a final step of regeneration in which morphogenesis is already strongly engaged. In contrast, the relatively low number of up-regulated genes (2619) between the last step of regeneration and the non-amputated control reveals the fact that at 5 dpa, the transcriptional program of regeneration is almost over.

Interestingly, we also observed very important numbers of down-regulated genes during the course of regeneration. This is notably the case when comparing 0 to 1 dpa and 1 to 2 dpa conditions where more than 6000 genes are down-regulated. The maximum of down-DEG is observed between 2 and 3 dpa stages (n> 9500). 2dpa (or stage 2) corresponds to the early blastema formation, when the cells are the less differentiated, while at 3 dpa blastema cell differentiation is well initiated, with notably the initiation of nervous system structures as well as muscles reformation.

Those down-DEG observed between this transition could correspond to the genes involved in the formation of a less differentiated structure at 2 dpa. Altogether those results highlight again the dynamic nature of the transcriptional landscape between early and late blastema stages. The lowest numbers of down-regulated genes are found comparing 3 dpa to 5 dpa and 5 dpa to NA. As for the up-DEG, those relatively low numbers reveal the fact that at 5 dpa, the transcriptional program of regeneration is almost over.

In addition, we have generated volcano plots to represent DEGs between sequential conditions (Fig. 3B). As already noticed from the heatmaps (Fig. 3A), comparisons of early stages (0 to 1 dpa and 1 to 2 dpa) show comparable numbers of up and down genes, with similar proportions of up- and down-regulated genes (and comparable significance levels). In contrast, comparisons between 2 to 3 dpa and 3 to 5 dpa show very different situations between up- or down-regulated genes. As mentioned before, many genes are down-regulated at 3 dpa in comparison to 2 dpa, which is strikingly visible in the volcano plot. Conversely, many genes are up-regulated at 5 dpa in comparison to 3 dpa. The late steps of regeneration also harbor comparable up- and down-regulated genes. Altogether, these results highlight the fact that the regeneration process is highly dynamic at the transcriptional level, and that the blastema stage (3 dpa), is very distinct from the other regeneration stages we sampled. It is worth to note that we included in our samples the regenerated structure plus the five anterior segments abutting the amputation plane. With this strategy, we aimed to obtain potential transcriptomic information triggered by the amputation signal coming from non-amputated tissues. Indeed, we previously reported that dedifferentiation processes of tissues abutting the amputation plans are potentially at play during blastema formation (41, 55). The risk of such strategy, however, is to buffer subtle differences between samples, with the transcriptional information from the five segments. However, the large numbers of DEGs (up- or down-regulated) identified in all the comparisons strongly indicate that we captured an important part of the transcriptional dynamic of posterior regeneration. Next, we examined the enrichment of Gene Ontology (GO) terms (over-representation analysis, focus on Biological Processes), of the DEG lists, for each comparison (Figure 4 and Additional file 14). For many comparisons (0 *vs* 1dpa, 0 *vs* 3dpa, 1 *vs* 2dpa, 1 *vs* 3 dpa, 1 *vs* NA, 2 *vs* 3dpa, 2 *vs* NA, 3 *vs* 5 dpa, 3 *vs* NA), the enrichment in GO term highlights the importance of ribosomal and mRNA biogenesis and processes for regeneration. This could be related to that fact that for those comparisons, impressive numbers of genes are dynamically expressed (from about 9000 to 16 000 DEGs (up- and down cumulated), depending on the conditions considered). The 0 *vs* 2 dpa comparison, in contrast, shows the importance of terms related broadly to morphogenesis (*e.g.* sensory, digestive, circulatory system), as well as extracellular organization. The 1 *vs* 5 dpa comparison, as well as the 5 dpa *vs* NA, in addition to some morphogenesis terms, reveal the importance of metabolic processes during these regeneration stages.

**Figure 4:**
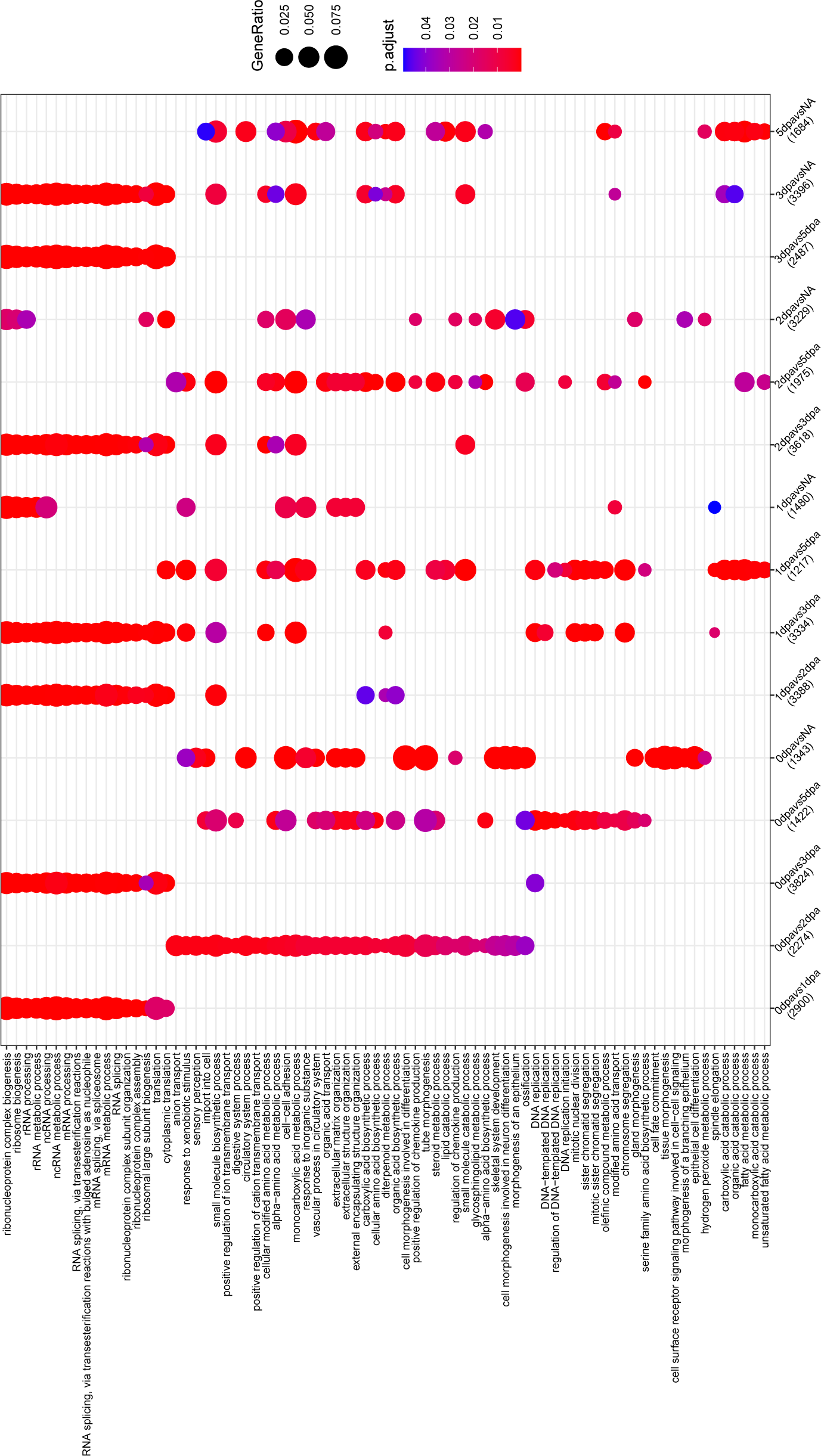
Gene ontology enrichment analysis of differentially expressed genes (DEGs) across *Platynereis dumerilii*’s regeneration. DotPlot showing most significant (Top 10) common over-represented Biological Process GO terms among each DEGs comparison (represented in columns). Circles area is proportional to the fraction of transcripts in each comparison (columns) falling into the corresponding GO term (lines), colors indicate the p-value of the enrichment.

#### • Identification of condition-specific differentially expressed genes and key gene categories

In addition to this very broad GO terms analysis, we aspired to gain more detailed insights into the regeneration transcriptional landscape.

To this end, we) identified the specific DEG between sequential conditions (Figure 5, Additional file 15); ii) identified unique DEG significantly specific to one stage (Additional file 16); iii) focused on genes involved in regulation (*e.g.* transcription factors and RNA-binding proteins, Additional file 17) and iv) specifically investigated nervous system and stem cells genes markers (Additional files 17 and 18).

**Figure 5:**
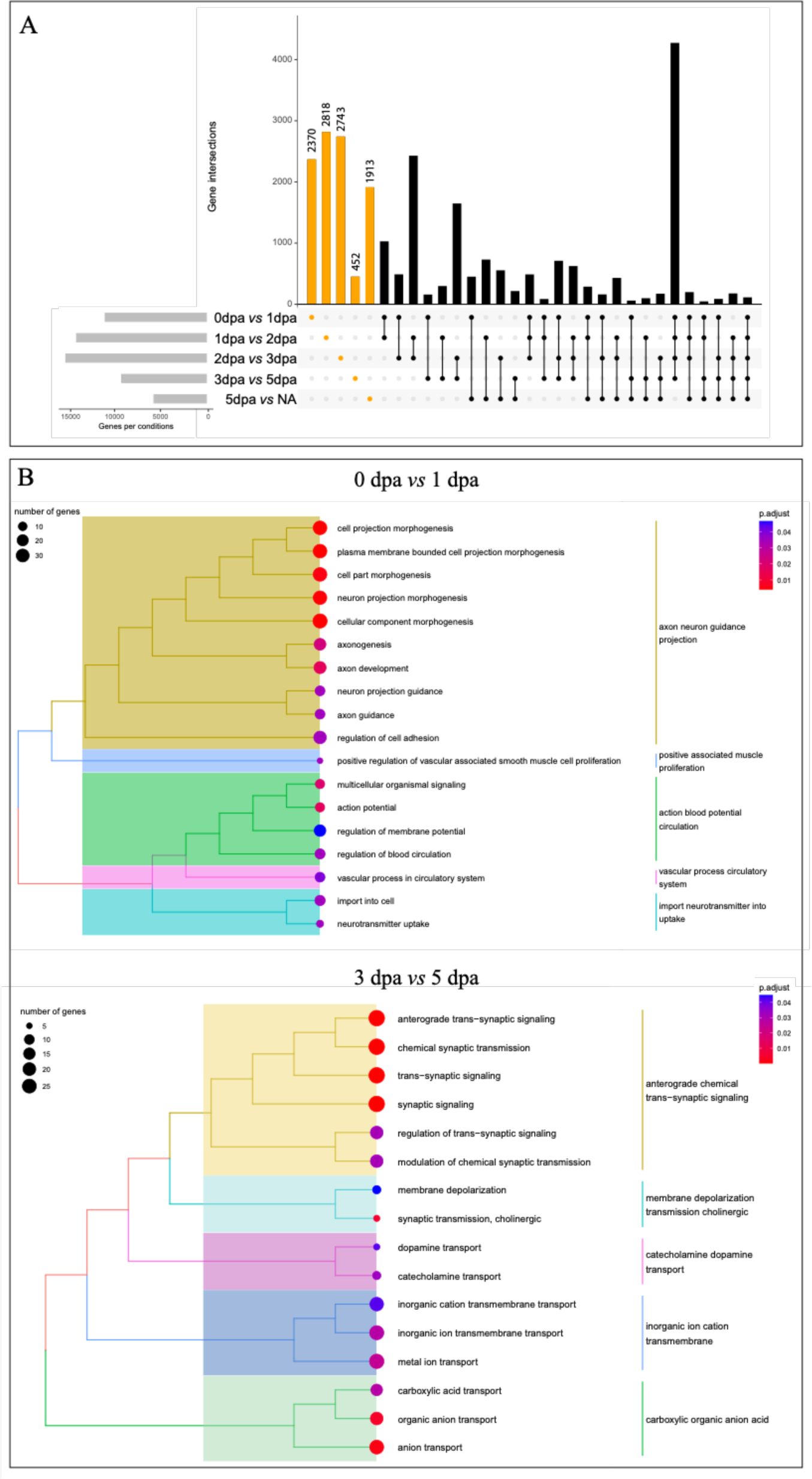
Comparison of gene expression between sequential regeneration stages. A) UpSetR plot showing the number of DEGs (differentially expressed genes) that are shared between comparisons (e.g. 2370 genes are identified as DEGs between 0 dpa and 1 dpa). Only sequential comparisons were considered (0dpa vs 1dpa; 1dpa vs 2dpa and so on). Orange bars represent DEGs unique to one comparison, black bars represent DEGs identified in more than one comparison. B) Treeplot representation of clustered GO term enrichment analysis (Biological Processes category) for two major sequential comparisons, namely 1 dpa vs 1 dpa, and 3 dpa vs 5 dpa. Circles area are proportional to the number of genes in each annotation, the colors represent the p-value of the over-representation.

First, we identified, quantified and analyzed DEGs specific to a sequential comparison between two stages (Fig 5). We identified 2370 DEG that are specific to the comparison 0 *vs* 1 dpa (or more specifically between the immediate amputation stage and the wound healing stage, Fig 5A). A GO term analysis (Biological Processes) revealed an involvement of neuronal projection guidance and the requirement of blood circulation system (Fig 5B). For the next comparisons, 1 to 2 dpa and 2 to 3 dpa, we identified 2818 and 2743 DEGs, respectively (Fig 5A). However, a GO term analysis did not highlight key elements. In contrast, when comparing 3 to 5 dpa (*i.e.* the blastema stage *versus* the differentiating regenerating structure), we identified a smaller number of DEGs (452) (Fig 5A). Nevertheless, a GO term analysis suggest a role of nervous system signaling, more specifically synaptic signaling, as well as cholinergic signaling, and dopamine transports (Fig 5B, Additional file 15). These results uncover major elements regarding the very early step of regeneration. First, the enrichment of GO terms related to blood and vascular circulation most likely corresponds to the immediate response to the amputation: containing and stopping the blood flow generated by the dorsal and ventral vessels opening following injury. Second, the nervous system appears as a meaningful component of both very early steps of regeneration but also later on during morphogenesis. Nerves and neural signaling have been identified as key players for regeneration in both vertebrates and non-vertebrates animals since centuries (56), including annelids (17).

This so-called nerve-dependency of regenerative processes can be of various types. Nerves can be direct sources of local signals, and while this type of nerve-dependency has never been evidenced so far in annelids (56), the major role of axon projections between 0 and 1 dpa is worth to note. We previously evidenced the appearance of a nerve ring underlying the wound epithelium (at 1 dpa), coming from the ventral nerve chord (41) supporting this hypothesis and calling for further investigation of this question in the future. The later role of nervous system during blastema morphogenesis (3 to 5 dpa) corresponds to the differentiation of neural progenitors of both the VNC and the peripheral nervous system of the developing segments, as evidenced previously thanks to the expression patterns of various neural markers (41). Those results call for a better functional understanding of the nervous system during regeneration in *Platynereis*.

Next, we identified unique DEGs, *i.e.* genes that were significantly up-regulated in only one stage compared to all other stages (Additional file 16). In most stages, those unique genes are in low numbers, except for the 2 dpa stage (n=164, 169, 624, 126, 13 and 121 for 0, 1, 2, 3, 5dpa and NA respectively). Looking at the functional annotation of these genes highlighted crucial features for regeneration. For stage 0, we found many genes related to immunity and inflammatory responses (Macrophage mannose receptors, Toll – like receptors, etc. – Additional file 16) as well as response to external stimulus. The specific expression of these genes at this very early stage is indicative of either an immediate transcriptomic response to injury from the immune system or an accumulation of cells from the immune system to the injury that would be enriched in the sampled tissues. Later on, at 1 dpa, other genes related to the immune system are also specifically expressed but to a lesser extent. In addition, we could identify genes related to the cytoskeleton, as well as development. Among developmental genes, the gene *atonal,* a member of the bHLH family (57) is expressed specifically at the stage 1. *Atonal* is a marker of peripheral sensory neurons during larval development in *Platynereis*, under the control of the BMP signaling pathway (58). Its function during regeneration remains to be discovered, especially at such early stage. The stage 2 is characterized by genes related to epigenetic mechanisms, notably chromatin remodeling and histone modifications. Epigenetic processes, more specifically DNA methylation, have been shown to be of major importance during *Platynereis* larval development as well as early stages of regeneration (55). Indeed, the use of an hypomethylating agent strongly delays the regeneration process that is blocked as early as stage 2.

At 2 dpa, we also found the presence of another major member of nervous system specification: *Pax2/5/8*. This transcription factor has been proposed to specify dorsal interneurons in *Platynereis* larvae (58). Its presence at this stage of *Platynereis* regeneration highlights, again, the potentially major role of nervous system signaling during the initiation of regeneration. At 3 dpa, genes specific to this blastema stage are mostly related to muscle morphogenesis, while at 5 dpa, no major biological process emerged from GO term analysis. Finally, the non-amputated stage is characterized by the presence of members of signaling pathways, notably Wnt. Among them are the ligand *Wnt6* and a Frizzled family gene modulating Wnt signaling, *Sfrp3/4* (59–61). At 48 hours post fertilization (hpf) larvae, *Wnt6* is expressed in the pygidium and proctodeal area, suggesting a role in the patterning of this region (61), that could be also at play during posterior growth.

Next, we decided to apply a strategy akin to a gene-candidate approach to specifically assess the importance of major gene expression regulators during regeneration: transcription factors (TF) and RNA-Binding Proteins (RBP) (Additional file 17). TFs, by controlling the rate of gene transcription, are of uttermost importance during morphogenetic processes and notably regeneration (62). RBPs are involved in many post-transcriptional processes leading to a fine-scale regulation of mRNA stability, localization and translation during development in animals (63, 64). The rationale here is to identify regeneration stages in which these types of regulators could be specifically enriched. To this end, we used a comprehensive animal TF database (AnimalTFDB 4.0) that contains 1611 TF from mouse (65), and identify potentially homologous genes in our transcripts (based on BLASTX analyses). We annotated 5248 of our transcripts as TFs, and among them, those that are specifically DEGs between sequential conditions of regeneration (88 ≤ n ≤ 236, Additional file 17). Using hypergeometric tests, we tested whether TFs are over-represented among these DEGs (in sequential comparisons). It appears that TFs are not over-represented for none of the sequential conditions (all p-values = 1). In contrast, when applying the same approach for RBPs (using the RBPDB database that contain 405 RBP from mouse (66)), we could highlight an over-representation of these gene expression regulators category for all conditions excepted one (5dpa *vs* NA) (Additional file 17). RBPs are thus especially at play during both the initiation of regeneration, and the blastema formation and differentiation, while it is no longer the case during the last step of growth and differentiation. These interesting observations are in line with previous experimental results showing the expression of many RBPs, such as *Pdu-piwi* or *Pdu-vasa,* during posterior regeneration (and elongation) in *Platynereis* (40, 41). Those genes are known to be expressed in somatic and/or germinal stem cells in many organisms (40, 67–69) and are thus considered as “stem cells” markers, forming what has been named the “Germline Multipotency Program” or GMP (70).

As mentioned before, the role of the nervous system and neural signaling is instrumental during regeneration of many metazoan species (56). In addition, stem cells, while not being the only source of cells in various regeneration contexts, play indisputably a major role in blastema formation during many regeneration model systems (*e.g.* planarians, cnidarians, acoels, etc. (71–73)). We thus specifically investigated nervous system and stem cells genes markers by two different means in our dataset. First, applying the same methodology as for TFs and RBPs, we used specific nervous system and stem cells related databases (74) that we compared to our DEGs associated to step-wise comparisons. While nervous-system related genes are not enriched in any conditions, SC genes are, for three comparisons (namely 1 *vs* 2 dpa, 2 *vs* 3 dpa and 5 dpa *vs* NA, Additional file 17). In addition, we also explored in detail a specific selection of “nervous system” (NS) and “stem cells” (SC) genes (n= 22 and 32, respectively) that were already well known and annotated in *Platynereis* as larval development or regeneration genes (Additional file 18) (40, 57, 58, 75, 76). Regarding our selection of NS genes, we observed that many of them are up regulated at 3 dpa in comparison to 2 dpa (*e.g. Sox B1, Neurogenin (ngn), Olig, Ash1, Pax 2/5/8*) (Additional file 18). Among them, *SoxB1, Olig, Ngn* and *Ash1* are part of the ancestral bilaterian toolkit of neural developmental genes (57, 76). Their role in patterning larval central nervous system has been well established in *Platynereis.* The activation of this nervous system program at 3 dpa coincides with the time at which the nervous system is being reformed during regeneration (41), suggesting that NS formation during both regeneration and development could rely on a similar transcriptional landscape. Within the “stem cells” genes, we mainly focused on the GMP signature previously mentioned and identified *Ago, Pumilio, PufA, Smb, Tudor2, Brat, Gustavus, Ap2* and *Lin28* as being up-regulated at 1 dpa in comparison to 0 dpa (Additional file 18). These results, suggesting an up-regulation of this multi/pluripotency program at the wound-healing stage are in agreement with previous experimental data. Indeed, various GMP members are intensely expressed in the wound-epithelium during *Platynereis* posterior regeneration, including for instance *Ap2* (41). Altogether, these elements support the hypothesis that blastema formation in *Platynereis* may rely on dedifferentiation events, as well as the fact that amputation potentially induces extensive reprogramming of differentiated cells into progenitor cells (41).

Overall, these extensive analyses of differentially expressed genes during various regenerations stages unveil important contributions of the nervous system and RBPs for different steps of regeneration. Many additional elements, especially members of key signaling pathways are also active during these stages and deserve a more in-depth investigation; they will be the subject of further studies.

#### • Co-expression clusters during posterior regeneration

With the aim to better assess and visualize the global transcriptional landscape of posterior regeneration, we performed a clustering analysis to identify differentially expressed genes sharing a similar expression dynamic (Figure 6). 12 clusters were defined and their dynamics were represented by both a heatmap (Figure 6A) and specific plots (Figure 6B). Among them, three clusters (clusters 3, 4 and 5) display a dynamic of gene expression that is upregulated at stage 1. 4077 genes are upregulated at 2 dpa (cluster 6), while cluster 10 reveals 2434 genes downregulated at this stage. 5415 genes have their highest expression at stage 3 (cluster 7). Clusters 8 (634 genes) and 9 (3091 genes) show the upregulations of genes at stage 5.

**Figure 6:**
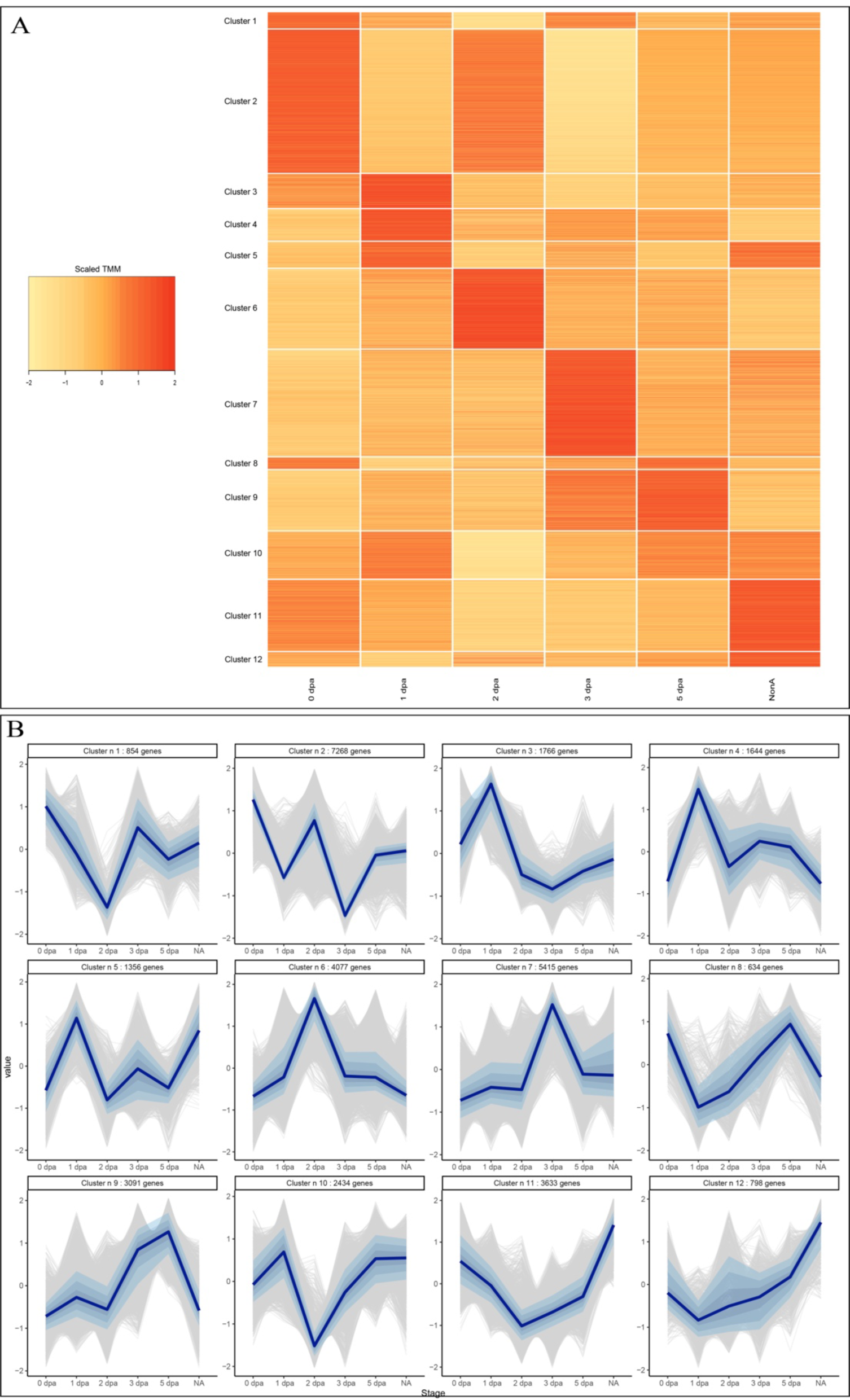
Clustering of genes based on their expression dynamics during regeneration in *Platynereis dumerilii*. A) Heatmap representation of the expression of each transcript sorted per cluster (n=12) throughout the posterior regeneration. Each line represents a transcript within its cluster, each column the regeneration stage considered. Colors indicate expression values. Expression values were taken as scaled TMM values (i.e. scaled down to have a mean of 0 and a standard deviation of 1). B) Gene expression profiles for each individual cluster (n=12). The number of transcripts contained in each cluster is indicated above each profile. Gray lines represent expression profiles of individual genes, dark blue lines represent the median expression across all genes in the cluster. Shaded blue areas around the median represent values in which 20%, 40% and 60% of expression values centered around the median are contained, respectively. All individual genes, for each of the 12 clusters, are listed in Additional file 20.

As previously done for DEGs, we identified clusters specifically enriched in TFs and RBPs, as well as NS and SC genes, using hypergeometric tests (Additional file 17). We did not find any cluster enriched in TF or SC genes, however clusters 2 and 6, that contain collectively more than 11000 genes upregulated at stage 2, harbor a specific enrichment for RBPs. These results are coherent with the previous tests made that showed an over-representation of RBPs during blastema formation, as well as already mentioned experimental results (41). In addition, cluster 7 – containing genes with the highest expression at stage 3, as well as cluster 11 (upregulation at 0 and NA stages) show an enrichment for NS genes. Our previous hypergeometric tests did not highlight a specific comparison between regeneration stages with an NS enrichment. However, previous experimental data demonstrated that the nervous system, and more specifically the ventral neurectoderm is specified at stage 3 (41, 77), which is in line with those results.

For each cluster, we also identified which GO terms were enriched (with a focus on biological processes - Additional file 19) and when appealing results were found, we investigated the genes content of those clusters. No significant enriched GO terms were obtained for clusters 3, 5, 6, 12. However, we found particularly interesting elements for clusters 7, 9, and 11.

The cluster 7, that reveals an upregulation of genes expression at stage 3, has an enrichment of GO terms related to morphogenetic processes and signaling (Additional file 19).

More precise investigation of the genes list for this cluster 7 revealed the presence of genes related to circulatory and blood system (*e.g. thrombospondin*), nervous system (such as the midline markers *Slit* and *Netrin,* see below (41, 58)), as well as many members of various signaling pathways. Among them, cluster 7 contains key members of the BMP signaling pathway: a *BMP-like* receptor, as well as its antagonist *Chordin*. In *Platynereis* larvae, BMP signaling is involved in the nervous system formation (58), while a similar function during regeneration remains to be evidenced, identifying key BMP members in the cluster 7 could be *a priori* compatible with such function. Transcription factors encoded by *Hox* genes, more specifically *Hox1* and *Hox4* are also found in this cluster 7 and they both show expression in nerve cells during *Platynereis* regeneration (in the VNC and VNC plus peripheral ganglia, respectively) (77). Cluster 7 also comprises the transcription factor *Gli* and its transmembrane activator *Smoothened,* both being key members of the Hedgehog (Hh) pathway, previously shown to regulate segments formation in *Platynereis’* larvae (78). Again, no information regarding Hh pathway during posterior regeneration in annelid is available so far, but a role in segmentation, akin to what is known for larvae, is conceivable. Two homeobox genes of the NK family (*i.e. NK1, NK4*), also present within the cluster 7, are involved in segment formation as well (79). Other signaling pathway components (*e.g.* some *Wnt* ligands and *Frizzled* receptors, as well as *Hes* and *Delta-like* genes, (60, 80–82)) are found in this cluster. Altogether those results are in line with the fact that segment formation as well as nervous system reformation occur at stage 3, as experimentally established previously (41).

The cluster 9, that reveals an upregulation of genes expression at stages 3 and 5, has an enrichment of GO terms mostly related to cell cycle (Additional file 19). Thus, exploration of cluster 9 genes highlights the presence of well-characterized cell cycle genes, including *Cyclin B1* and *PCNA*, intensely expressed from stage 3 onward during *Platynereis* regeneration. We previously experimentally evidenced, through EdU labelling, *in situ* hybridization, as well as cell proliferation inhibitors that an intense phase of cell proliferation is mandatory from stage 3 onward, for a proper regeneration process (41). Those RNA-seq data are thus in agreement with experimental data. In addition, some signaling pathways members (*e.g. Hes* genes, *Fzd9/10* receptor, *Wnt16* ligand) are also found in this cluster 9.

We then concentrated our interest on the cluster 11 that may reveal important genes for late stage of regeneration. Cluster 11 is characterized by an enrichment of GO terms related to developmental (notably epithelium and circulatory system) and growth processes, and more specifically, the MAPK pathway (Additional file 19). Indeed, *Maf* and *c-jun* transcription factors, as well as various members of the Mitogen-activated protein kinase (MAPK) pathway are found in this cluster. Others elements associated with Notch, FGF, Hh, BMP, WNT, and potentially Toll and NFkB pathways could also be evidenced. Among and above them, some genes are related to proctodeum patterning (*i.e. Wnt7* and *Wnt10*) or segmentation (*Wnt10, Wnt16, Nk1, Lbx,* (60, 79).

Altogether these fine-scale explorations of cluster contents bring to light, non-unexpectedly, the major involvement of many signaling pathways during *Platynereis* posterior regeneration. Among them, the Wnt pathway is an obvious interesting candidate to investigate in the future, due to its implication in regeneration processes at the metazoan level (including annelids such as *Syllis gracilis* (37) and *Eisenia andrei* (34)), as discussed previously (2). Beyond Wnt, other major signaling pathways (notably BMP, Notch, MAPK, and Hh) have been shown to be active in regenerative processes in a lower number of species so far. These pathways would be worth investigating in the future. For many of them, their functions during *Platynereis* larval development were already assessed (58, 78, 80, 82–85) and establishing their roles during regeneration would bring new key data to clarify the extent of recapitulation of development during regeneration in animals. Importantly, these studies will allow to apprehend the fine regulation of regeneration processes in an annelid species, which is a mandatory pre-requisite to assess the evolutionary history of regeneration at the metazoan scale.

With the objective to demonstrate the quality and validate the presented RNA-seq dataset, we performed whole mount *in situ* hybridizations (WMISH) on seven genes pertaining to three of those clusters (6, 7 and 11) (Figure 7). These expression patterns were determined during the whole course of regeneration (7 stages), from 0 to 5 dpa, also encompassing the stage 4 (or 4 dpa, not sampled for the transcriptomic analyses). Of note, as WMISH are not working efficiently on non-amputated worms, we used 15 dpa worms, as a proxy of a non-amputated stage, as previously explained (40).

**Figure 7:**
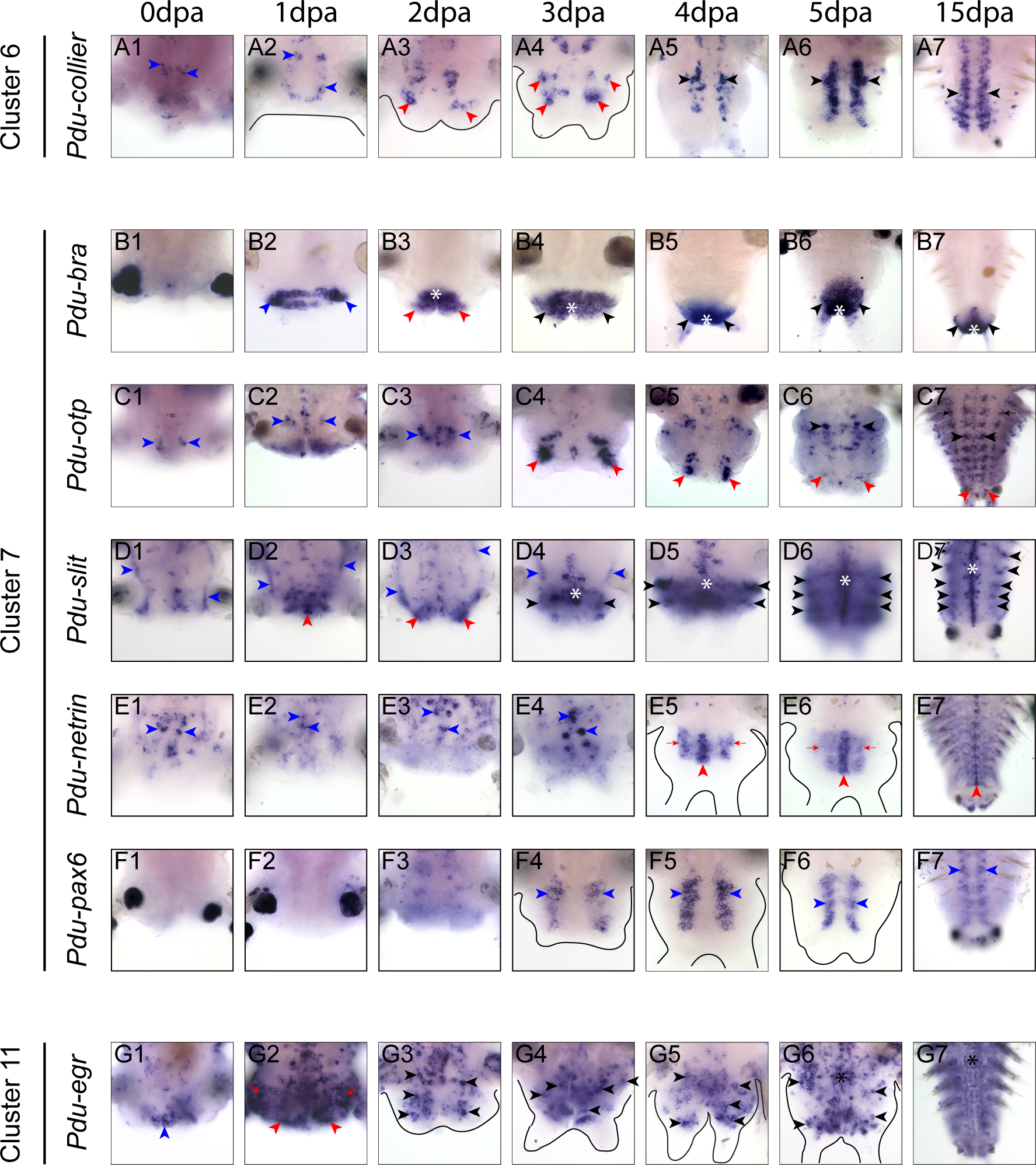
Expression analysis of a selection of genes representatives of clusters 6, 7 and 11 during Platynereis’ regeneration. Whole-mount in situ hybridizations (WMISH) for the genes whose name is indicated are shown for all regeneration stages (0 dpa to 5 dpa) as well as 15 dpa as a proxy of non-amputated control. All panels are ventral views (anterior is up). Outlines of the regenerated structures are depicted in black for the most transparent samples. A) WMISH for Pdu-collier. Blue arrow heads: differentiated neurons of the ventral nerve chord (VNC); red arrow heads: neurons present inside the blastema; black arrow heads: neurons inside the reformed VNC. B) WMISH for Pdu-bra. Blue arrow heads: wound epithelium. Red arrow heads: small blastema. White asterisks: anus. Black arrow heads: reforming/reformed pygidium. C) WMISH for Pdu-otp. Blue arrow heads: VNC of non-amputated segments. Black arrow heads: VNC of regenerated structures. Black arrows: peripheral structures. Red arrow heads: basis of the anal cirri. D) WMISH for Pdu-slit. Blue arrow heads: enteric nerve net. Red arrow heads: Mass of cells expressing slit at early stages of regeneration. White asterisk: midline. Black arrow heads: ectodermal stripes. E) WMISH for Pdu-netrin. Blue arrow heads: scattered cells inside the midline of non-amputated segments. Red arrow heads: midline of regenerating structures. Red arrows: midline-contiguous mediolateral domains expressing netrin. F) WMISH for Pdu-pax6. Blue arrow heads: longitudinal ventro-lateral lines of cells. G) WMISH for Pdu-egr. Blue arrow heads: mass of ventral abutting the amputation plane. Red arrow heads: wound epithelium. Red arrows: strong expression of egr in the segment abutting the amputation plane. Black arrow heads: scattered egr-expressing cells, potentially neurons. Black asterisk: VNC.

Regarding the cluster 6 (expression peak at 2 dpa), we selected the gene *Pdu-Collier*. This gene encodes a Helix-Loop-Helix transcription factor known to be mainly involved in the ventral nerve chord formation in *Platynereis* larvae, being more specifically expressed in post-mitotic neural cells (75). At 0 and 1 dpa, *Pdu-Collier* is expressed only in few specific differentiated neurons of the ventral nerve chord (VNC) of the non-amputated segments (above the amputation plane) (Fig. 7 A_1_ and A_2_, blue arrow heads). At 2 dpa, few neurons are present in the small blastema and, their number drastically increased at 3 dpa (Fig. 7 A_3_ and A_4_, red arrow heads). At 4, 5 and 15 dpa/NA, more neurons are expressing *Pdu-Collier* in two longitudinal rows that correspond to the VNC (Fig. 7 A_5_ to A_7_, black arrow head).

We also investigate the expression patterns of 5 genes (*Pdu-Brachyury, Pdu-Otp, Pdu-Slit, Pdu-Netrin, Pdu-Pax6*) corresponding to the biggest cluster (number 7, peak of expression at 3 dpa). *Pdu-Brachyury* (*Pdu-Bra,* a T-box transcription factor) is a known marker of posterior structure, notably the pygidium and anus during *Platynereis* posterior elongation (86). Our expression patterns during the time course of regeneration are in agreement with this fact.

Excepted at 0 dpa, where no expression is detected (Fig. 7 B_1_), *Pdu-Bra* is found exclusively in posterior tissues, in the wound epithelium at 1 dpa (Fig. 7 B_2_, blue arrow heads) in the anus and partly in the small blastema at 2 dpa (Fig. 7 B_3_, red arrow heads) in the (future) pygidium and anus from 3 to 15 dpa/NA (Fig. 7 B_4_ to B_7_, asterisks and black arrow heads respectively). We then determine the expression patterns of the TF *orthopedia* or *Pdu-Otp* a conserved marker of sensory cell in larvae (87). During regeneration *Pdu-Otp* is expressed in various territories: (i) in the VNC of non-amputated segments at 0, 1 and 2 dpa (Fig. 7 C_1_ to C_3_, blue arrow heads), (ii) in the VNC at 5 and 15 dpa/NA (Fig. 7 C_6_ to C_7_, black arrow heads), (iii) at the basis of the anal cirri from 3 dpa onward (Fig. 7 C_4_ to C_7_, red arrow heads), (iv) in peripheral structures, potentially the peripheral nervous system, in very differentiated segments (Fig. 7 C_7_, black arrows). Next, *Pdu-Slit,* an established marker of ventral midline during both larval development and regeneration was examined (41, 58). During regeneration, *Pdu-Slit* is found in different locations in addition to the midline of both the non-amputated segment and regenerated parts (Fig. 7 D). From 0 to 3 dpa, an interesting expression in the gut epithelium lining is also observed, potentially in the nerve net encompassing the gut (Fig. 7 D_1_ to D_4_, blue arrow heads). At 1 dpa, a large mass of cells abutting the amputated VNC starts to express *Pdu-Slit,* this expression extends to the whole small blastema at 2 dpa (red arrow heads, Fig 7 D_2_ to D_3_). From 3 to 15 dpa/NA, as previously reported (41), *Pdu-Slit* is expressed in the ventral midline cells (white asterisks), as well as in ectodermal segmental stripes (Fig. 7 D_4_ to D_7_, black arrowheads). Expression patterns of a second marker of midline, *Pdu-Netrin* (58), were also determined (Fig. 7 E). From 0 to 2 dpa *Pdu-Netrin* is expressed in few scattered midline cells of non-amputated segments, and no expression is found in the regenerated parts (Fig. 7 E_1_ to E_3_, blue arrow heads). At 3 and 5 dpa, strong expression in midline cells (red arrow head) is observed as well as in two mediolateral domains (red arrows) abutting the midline (Fig. 7 E_4_ to E_6_). Those medio-lateral expressions progressively fade away at 5 dpa. At 15 dpa, expression pattern in the midline is maintained while very restricted to few cells (Fig. 7 E_7_). A potential expression in parapods is also observed (Fig. 7 E_7_). Lastly, we looked at *Pdu-Pax6*, a gene expressed in the ventro-lateral part of the trunk ectoderm in both larvae and regenerating structure (41, 58) (Fig. 7 F). As expected, *Pdu-pax6* is expressed from 3–15 dpa/NA in two longitudinal ventro-lateral bands of cells (Fig. 7 F_4_ to F_7_, blue arrow heads) in the anterior part of the regenerated region, which likely correspond to neurectodermal domains of the future segments. No expression is observed at previous stages.

Regarding the cluster 11 (that reveals two expression peaks of at 0 and 15 dpa/NA), we focused our interest on a key regeneration gene, the transcription factor encoding-gene *Egr* (for early growth response) (Fig. 7 G). This gene has been identified as regulatory master gene for the initiation of regeneration in various model species such as the acoel, *Hofstenia miamia* (71), the sea star *Patiria miniata* (16), the planarian *Schmidtea mediterranea* (88) and more recently the annelid *Eisenia andrei* (34). *Pdu-Egr* starts to be expressed at 0 dpa, in a mass of cells, abutting the amputation plan, in the ventral part (Fig. 7 G_1_, blue arrow head). At 1 dpa, its expression is intense in the wound epithelium (red arrow heads) as well as all the non-amputated segment (red arrows) abutting the amputation plane (Fig. 7 G_2_). From 2 to 5 dpa, *Pdu-Egr* has a salt and pepper expression in the ventral part of the regenerated structures (black arrow heads), potentially in some neurons, as well as the VNC (black asterisk) at 5 dpa (Fig. 7 G_3_ to G_6_). At 15 dpa/NA stage, *Pdu-Egr* is mainly expressed in some cells of the VNC (Fig. 7 G_7_).

Altogether, these expression patterns from 7 genes that are members of the clusters 6, 7 and 11 highlight their transcriptional dynamics during the course of *Platynereis*’ posterior regeneration. It is worth to note that not all *of them* (*e.g. Pdu-Collier*) are always perfectly congruent with their expected dynamics based on the cluster they are assigned to. This can be due to two reasons. First, among the genes that are considered to be part of one cluster, not all follow the general tendency of this cluster (see the grey lines *versus* blue ones in Figure 6). More importantly, our transcriptomic data were done on samples encompassing both the regenerated structure as well as the 5 previous non-amputated segments. Because of this, the gene expression signal we will collect with our RNA-seq data will be a mix of those two structures, which can blur the signal we can obtain. Even with this slight limitation, those RNA-seq data covering the course of posterior regeneration in *Platynereis* offer a unique opportunity to dissect the precise transcriptional landscape of major steps of this process and give important insights for future specific studies.

### Conclusions

Restorative regeneration is a fundamental process shared across many animal lineages, yet its fine-scale understanding is still scarce for a majority of them. Elucidating how regeneration is regulated in a diversity of model species is a mandatory step to address key questions in the field, especially those regarding the evolution of this process in Metazoa. One way to approach this question of uttermost importance is to assess and compare the transcriptomic landscape patterning the regeneration of various structures. In this study, we provided a detailed characterization of the transcriptomic landscape of posterior regeneration in the annelid *Platynereis dumerilii*. This analysis highlighted the important contribution of the nervous system, as well as the major involvement of RBPs and signaling pathways during this process, all deserving mechanistic/functional exploration. In addition, we produced a high-quality Reference transcriptome for this front-line annelid model species that could serve as a valuable resource for both embryonic, larval and regeneration studies.

## Methods

### *Platynereis dumerilii*’s culture, biological material production, mRNA extractions, and sequencing

Individuals of *P. dumerilii* were obtained from the laboratory breeding culture established at the Institut Jacques Monod (France) (28). For all regeneration experiments, 3-4-month-old worms with 30-40 segments were used and amputations of the last 5-6 posterior-most segments were performed as previously described (28, 41). After amputation, worms were let to recover and regenerate in natural fresh sea water (NFSW) for the time necessary to reach the regeneration stage(s) of interest. Regenerating regions were then recovered in a tube on ice and immediately placed at -80°C. Three different series of biological samples were produced as follows (Additional file 1):

- Series 1: blastema tissues at 2- and 3-days post-amputation (stages 2 and 3). For each condition, 150 regenerating parts (and 3 biological replicates per stage independently produced) were recovered with as less as possible of non-amputated tissues (usually half a segment from abutting tissues).
- Series 2: 3dpa and non-amputated tissues. For 3dpa condition, 150 regenerating parts (with as less as possible of non-amputated tissue) as well as 150 non-amputated (NA) posterior parts (encompassing the pygidium, the growth zone and 5 segments) were recovered. No replicate performed.
- Series 3: 0dpa (= immediately after amputation, stage 0), 1dpa (stage 1), 2dpa (stage 2), 3dpa (stage 3), 5dpa (stage 5), as well as NA tissues during regeneration course. For each condition, 30 regenerating parts (or NA) plus the 5 abutting segments were recovered. Three biological replicates per condition were performed (Additional file 1).

For all samples, total RNA was extracted with the RNAqueous Total RNA Isolation Kit (Ambion) following manufacturer’s protocol. Total RNA quality was assessed using a Bioanalyzer (Agilent). Illumina mRNA-seq libraries were then prepared for samples series 1 and 3, by the sequencing platforms (Eurofins and Duke Institute for Genome Sciences and Policy, respectively).

For the samples series 2, library preparation and Nanopore sequencing were performed at the Ecole normale supérieure GenomiqueENS core facility (Paris, France). 10 ng of total RNA were amplified and converted to cDNA using SMART-Seq v4 Ultra Low Input RNA kit (Clontech). Afterwards an average of 10 fmol of amplified cDNA was used to prepare library following SQK-PBK004 kit (PCR Barcoding kit; ONT). After the PCR adaptater ligation a 0, 6X Agencourt Ampure XP beads clean-up was optimised and 2 fmol of the purified product was taken into PCR for amplification and barcodes addition with a 17 minutes elongation at each 18 cycles. Samples were pooled in equimolar quantities to obtain 36 fmol of cDNA and the rapid adaptater ligation step was performed. Libraries were multiplexed by 2 on a FLO-MIN106 flowcell according to the manufacturer’s protocol.

For the series 1 (Blastema tissues), Illumina deep sequencing with 150 bp paired-end reads was performed, while for the series 3 (regeneration timeline), 50 bp single-end reads were obtained, using Illumina HiSeq sequencing system. For the series 2, long reads were sequenced using the Oxford Nanopore Technology (ONT) and the MinION sequencing system (Additional file 2). More specifically, sequencing was performed with the SQK-PBK004 72-hour sequencing protocol run on the MinION MkIC, using the MinKNOW software (versions 20.10.6). A mean of 3.9 ± 0.6 million passing ONT quality filter reads was obtained for each of the 2 samples. Base-calling from read event data was performed by Guppy (version 4.4.2). All raw reads for individual sequencing libraries are deposited in the European Nucleotide Archive (ENA) under accession number detailed in the Additional file 2.

### Read processing and *de novo* transcriptomes assembly

All Illumina short raw reads were processed with Trimmomatic (89) using the following command to eliminate short and low-quality reads, as well as adapter sequences: ILLUMINACLIP:$TRIMMOMATIC_DIR/adapters/TruSeq3-PE.fa:2:30:10 SLIDINGWINDOW:4:5 LEADING:5 TRAILING:5 MINLEN:25 for paired-end reads and ILLUMINACLIP:$TRIMMOMATIC_DIR/adapters/TruSeq3-SE.fa:2:30:10 SLIDINGWINDOW:4:5 LEADING:5 TRAILING:5 MINLEN:25 for single-end reads.

ONT long raw reads were processed with TALC (90) with default options.

With the aim to produce a high-quality, as complete as possible reference transcriptome, we chose to combine several pre-existing and newly generated *de novo* transcriptomes. This strategy (Additional file 3) allows to include a diversity of biological datasets and to combine the advantages of several assembly pipelines and methods.

We performed a first assembly using only the blastema short reads (series 1) using Trinity (91) with default options and named it the Blastema transcriptome. We then carried out a second assembly using the blastema short reads (series 1) as well as the non-corrected ONT long reads (series 2), using rnaSPAdes (48) with default options. This assembly is named Hybrid 1 transcriptome. We then performed a third assembly, named Hybrid 2 transcriptome, also using the blastema short reads (series 1) and the ONT long reads (series 2), but using the Trinity suite. We finally assembled a reference transcriptome for *P. dumerilii* by combining these three newly generated transcriptomes (Blastema, Hybrid 1 and Hybrid2) with two others from the literature, namely from the adult head (45) and from developmental stages (44) using the EvidentialGene suite (92) with default parameters. The overall methodology of transcriptomes aggregation is depicted in Additional file 3. The order of fusion of transcriptomes has no influence on the result. The Reference *de novo* transcriptomes is deposited in the European Nucleotide Archive (ENA) under accession numbers detailed in the Additional file 2.

### Transcriptomes quality assessment

We assessed and compared the quality of the 6 transcriptomes (Blastema, Hybrid 1, Hybrid 2, Head, Embryonic, Reference) by computing a number of classic metrics. First, using Trinity (91), we computed total number of transcripts and genes, mean, min & max transcript length, transcript N50 (length of the shortest transcript for which longer and equal size transcripts cover at least 50% of the transcriptome sequences) (52), as well as Ex90-N50 (N50 for the 90% most expressed transcripts based on the blastema short reads) (Additional file 4). We further appraised the quality of the reads’ mapping on each transcriptome. For that purpose, the Blastema short reads (series 1) we generated, as well as the short reads used for the assembly of the head and embryonic transcriptomes (head and developmental stages dataset, respectively, see Additional file 2) were mapped on each transcriptomes using bowtie2 (93) with default parameters (Additional file 5).

We also assessed gene completeness for each transcriptome using BUSCO (Metazoan gene set) (Additional file 6) (94). Given the redundancy of transcriptome assemblies compared to genome assemblies, single-copy and duplicated genes were merged together. We finally evaluated how transcriptomes could recapitulate known coding and protein sequences. We computed the percentage of *P. dumerilii* coding sequences on the NCBI Nucleotide database (95) covered by at least one transcript and for that purpose; we blasted each transcriptome to the NCBI database using a e-value cutoff of 10^-5^ and reported the number of coding sequence with a match. Similarly, we computed the percentage of transcripts covered by at least one Swiss-Prot protein sequence (using the non-redundant database), using blast with e-value cutoff of 10^-5^ and the qcov output format to compute the query coverage (Additional file 6).

### Functional annotation of the *Platynereis* Reference transcriptome

ORF (open reading frame) predictions were performed with EvidentialGene. We annotated the Reference transcriptome using Trinotate (96) (Additional file 7). This tool generates for each predicted protein the Swiss-Prot top BLASTP hit, the PFAM domains, predictions of trans-membrane function (tmHMM), the eggnog annotation, the KEGG pathway, and the gene ontology (obtained with BLASTX, BLASTP & PFAM). We also performed BLASTX searches between the Reference transcriptome sequences and coding sequences of the following species (97): *Homo sapiens*, *Mus musculus*, *Danio rerio*, *Saccoglossus kowalevski, Drosophila melanogaster*, and *Aplysia californica* (Additional files 8 and 9). We computed the percentage of *P. dumerilii* sequences with a BLASTX hit in at least one species (e-value cutoff: 10^-3^, only the best hit was retained in case of multiple hits).

We finally performed additional BLAST analyses to assess the quality of our assembly and evaluate our previous BLAST results. We first used the transcript sequences of a well-reconstructed and annotated annelid genome, *Owenia fusiformis* (53), to perform BLASTX searches with the coding sequences of the 6 species mentioned above. Second, we performed a similar BLAST search, using our transcript sequences as indicated in (53), namely on a non-redundant gene database from well-annotated annelid genomes (*Capitela teleta*, *Dimorphilus gyrociliatus*, *Eisenia andrei*, *Lamellibrachia luymesi*, *Paraescarpia echinospica*, *Riftia pachyptila* and *Streblospio benedicti*). For both analyses, a e-value cutoff of 10^-3^ was used to consider a hit as significant.

### Gene expression levels and dynamic across *Platynereis dumerilii’s* posterior regeneration

#### Read processing and mapping

To study the dynamics of gene expression during regeneration in *P. dumerilii*, we used the Regeneration set of RNA-seq short reads (series 3, see above). Of note, these very short reads were not used to reconstruct the Reference *P. dumerilii* transcriptome.

After processing the reads (quality and adapter trimming, see above for details), we used kallisto (98) within the Trinity suite (91) to perform pseudo-mapping and quantification of the reads on the Reference transcriptome previously reconstructed (Additional file 11). After performing a PCA analysis, three samples were discarded as detected as outliers (one sample at 1dpa, 3dpa and 5dpa) (Additional file 10). We used edgeR (99) to process gene expression values. Read coverage per gene were processed to obtain TMM (trimmed mean of M values) (100) normalized expression data (Additional file 12). A gene was considered as expressed if its TMM values is superior or equal at 1.

#### Differential expression and clustering

We used edgeR (99) to identify differentially expressed genes in all possible comparisons (Additional file 13). Differentially expressed genes were defined as genes with a logFC value of either less than -1 or above 1, and an FDR value below 0.05. We used the UpSetR library in R (101) to quantify and visualize shared expressed genes between conditions and shared DEGs between comparisons, and the EnhancedVolcano library in R to build volcano plots (102).

We next identified DEG that are specific and unique to one stage. For that purpose, we determined upregulated DEG for a given stage in comparison to all the others (Additional file 16).

We then performed hierarchical clustering of DEG from median TMM values across replicates as expression values (median values were scaled to have a mean of 0 and standard deviation of 1) and Euclidian distances, using the hclust function in R (method “complete”). After visual inspection, we retained 12 clusters of co-expression that were reordered by expression patterns. A heatmap visualization of the clusters was performed using heatmap.2 from the gplots R package. The complete list of gene identifiers included in each cluster is available in Additional file 20.

#### Gene ontology and enrichment analysis for differentially expressed genes and clusters

We further analyzed the content of DEG lists as well as clusters. We performed Gene Ontology (GO) - term enrichment analyses using clusterProfiler (103). For the DEGs, a full list of enriched GO terms is provided in Additional file 14 and the top 10 for the category “Biological processes” are presented thanks to a dotplots. We specifically focused on the lists of genes generated from step-wise comparisons (*i.e.* 0dpa *vs* 1dpa, 1dpa *vs* 2dpa, 2dpa *vs* 3dpa, 3dpa *vs* 5dpa, 5dpa *vs* non-amputated). For clusters, GO terms enrichments were represented using the treeplot function from clusterProfiler, using “average” for the hclust method (Additional files 19 and 20). In addition, we computed over-representation of genes in specific “functional” categories, specifically transcription factors (TFs), RNA-binding proteins (RBP), as well as nervous system development (NS), and stem-cells markers (SC) thanks to available databases from the literature. For each group of genes (cluster or list of DEGs), we kept only *P. dumerilii* genes with a *M. musculus* homologous gene, and looked if that mouse gene was annotated as TFs (65), RBP (66), NS (based on gene list associated to nervous system GO terms), or SC (74) (Additional file 17). For each list of genes, we thus counted the number of genes matching each category. We computed over-representation p-values using hyper-geometric tests, correcting p-values for multiple testing (Benjamini-Hochberg FDR correction) (Additional file 17).

Finally, we focused our interest on selections of genes of peculiar interest in the context of regeneration. Hence, we manually computed lists of markers of stem cells and of the germline multipotency program (GMP, n=32) (40, 67, 70), as well as of neurogenesis and neural signaling (n=22, (58, 80)) (Additional file 18). We identified those genes in the Reference transcriptome, recovered their expression values, as well as the conditions where they are differentially expressed.

### Whole-mount *in situ* hybridization (WMISH) and imaging

WMISH on regeneration stages were performed as previously described (28, 41). Seven genes were selected: *Pdu-Collier* (PladumEVm010659), *Pdu-Brachyury* (PladumEVm013103), *Pdu-Otp* (PladumEVm027642), *Pdu-Slit* (PladumEVm001726), *Pdu-Netrin* (PladumEVm011161), *Pdu-Pax6* (PladumEVm031580), *Pdu-Egr* (PladumEVm012194). Bright field pictures for WMISH were taken on a Leica microscope DM5000B. Editing and compilation of Z projections were achieved using Adobe Photoshop CC 2017. The final figure panels were then compiled using Adobe Illustrator CC 2017.

## Additional files legends

### Additional file 1: Experimental design for the transcriptomic analysis of regeneration

Schematic representation of the five stages of regeneration (stage 0 to stage 5) and the non-amputated control, as well as their corresponding days post amputation (dpa, from 0 to 5) are shown. The amputation plan is represented by a red dotted-line on each schematic. From stage 0 to stage 5, the regenerated region (if any) as well as the 5 anterior segments were collected (around 30 individuals *per* stage) for the RNA-seq experiment. Experimental information is provided in blue on the right. In addition, main morphological and cellular steps during the course of regeneration are shown at the bottom.

### Additional file 2: Accession and detailed information about the datasets used in this study

Detailed information for each dataset includes: (i) information about the biological structure that was sampled (name, type and stage of the structure), (ii) the name of each samples, (iii) the data type (long read, short read, read length etc.), (iv) the permanent links to data and accession numbers, as well as (v) the reference.

### Additional file 3: Reference transcriptome assembly strategy

Schematic representation of the transcriptome assembly strategy followed in the study. We assembled the Reference by merging three transcriptomes generated in this study (Hybrid 1, Hybrid 2 and Blastema) that were assembled using different combinations of data and methods (see Methods and Results and Discussion), and two transcriptomes extracted from the literature: Head (45) and Embryonic (44). Merging transcriptomes was performed using EvidentialGene (92). The type of data used per transcriptome assembly is mentioned on the top line of the schematic, while the different assemblers are color-coded and their names mentioned at the bottom of the schematic.

### Additional file 4: Transcriptome assembly metrics

Spreadsheet 1: number of transcripts and genes in each of the six transcriptomes (when available)

Spreadsheet 2: detailed assembly metrics (*e.g.* number of transcripts with length above 10kb) for all six transcriptomes

Spreadsheet 3: ExXN50 values for all six transcriptome assemblies. Values represent the N50 (at least 50% of the assembled transcript nucleotides are found in transcripts longer than this value) for the X% top expressed transcripts. Each line corresponds to a percentage (*e.g.* 1 corresponds to the top 1% most expressed transcripts).

Spreadsheet 4: Diverse metrics for all six transcriptome assemblies. N50 (N40, N30, etc.) values represent the minimum length of transcripts in which 50% (resp. 40%, 30%, etc.) of all assembled nucleotides can be found as well as the median contig length and average contig.

### Additional file 5: Mapping statistics for all transcriptome assemblies

Each RNA-seq dataset (see Additional file 2) was mapped on all transcriptome assemblies. Numbers represent the percentage of reads that were mapped on each transcriptome. Metrics for each RNA-seq dataset are in individual spreadsheet.

### Additional file 6: Assembly completeness metrics

Spreadsheet 1: BUSCO completeness metrics for all transcriptomes were computed for Eukaryota, Metazoa and Arthropoda datasets. Categories “complete single-copy” and “complete duplicated” were merged. The total number of BUSCO groups searched is mentioned.

Spreadsheet 2: Number and percentage of previously annotated *P. dumerilii* transcripts (deposited in the NCBI database) matching each of the six transcriptomes.

Spreadsheet 3: Number and percentage of transcripts with a significant BLASTX hit on the non-redundant Swiss-Prot database for each of the six transcriptomes.

Spreadsheet 4: Distribution of the query coverage for the BLASTX on the non-redundant Swiss-Prot database for each of the six transcriptomes. For each transcript, the best non-redundant hit was taken and the percent of the query *P. dumerilii* transcript covered by the Swiss-Prot protein was computed. Numbers indicate the number and percentage of transcripts covered between 0 and 25%, between 25 and 50%, between 50 and 75%, and above 75%.

### Additional file 7: Trinotate annotation results for the *de novo* Reference transcriptome produced

Default trinotate output containing for each transcript various information (Pfam domain, SignalP, BLAST results …).

### Additional file 8: BLAST results for the Reference transcriptome

Top panel: Graphic representation of BLASTX and BLASTP results from Trinotate (see Additional file 7). Areas in red and blue represent the percentage of *P. dumerilii* transcripts with (respectively without) a significant BLAST hit.

Middle and bottom panels: Graphic representation of BLASTX results between our Reference transcriptome and six bilaterian species (*H. sapiens*, *M. musculus*, *D. rerio*, *S. kowalevski*, *A. californica*, *D. melanogaster*). Areas in green, orange and yellow represent the percentage of *P. dumerilii* transcripts with a significant BLAST hit on an annotated protein, with a significant BLAST hit on an uncharacterized protein, and without a significant blast hit (e-value cutoff = 10^-3^) respectively.

### Additional file 9: Reference transcriptome annotation *via* BLASTX results on 6 individual species

For each transcript, only the top significant BLASTX hit (and its annotation) is shown. Species considered are: *Mus musculus, Homo sapiens, Drosophila melanogaster, Aplysia californica, Saccoglossus kowalevskii* and *Danio rerio*.

### Additional file 10: PCA plot representing the relatedness of RNA-seq samples obtained for six regeneration stages

A) PCA with all samples, B) Outlying samples have been removed (see Main Text for details). Two to three replicates per stages are present. Each dot represents an RNA-seq sample. Colors represent biological conditions (*i.e.* regeneration stages and non-amputated worms, indicated on the right), biological replicates are also indicated (by a number on the name of the samples).

### Additional file 11: Raw read count for the regeneration RNA-seq dataset

Expression value for all transcripts for each RNA-seq sample are shown. Outlying samples have been removed.

### Additional file 12: TMM expression values for the regeneration RNA-seq dataset

Expression values for all transcripts for each RNA-seq sample are shown. Outlying samples have been removed.

### Additional file 13: List of differentially expressed genes (DEGs) for all comparisons of regeneration stages

A gene is considered as differentially expressed with an FDR < 0.05 and a logFC > 1 or < -1. The table includes various annotations (BLAST).

### Additional file 14: Significantly over-represented GO terms for all lists of DEGs (adjusted p-value < 0.05)

Only Biological processes GO terms are shown, for all sequential comparisons of regenerations stages presented in each spreadsheet.

### Additional file 15: Significantly over-represented GO terms for DEGs specific to sequential comparisons

DEGs specific to a given sequential comparison are, for instance, genes differentially expressed in 0 dpa *vs* 1 dpa but not differentially expressed in 1 dpa *vs* 2 dpa, 2 dpa *vs* 3 dpa, 3 dpa *vs* 5 dpa or 5 dpa *vs* NA. Those genes are considered as unique for the 0 dpa *vs* 1 dpa comparison. Only Biological processes GO terms are shown for all DEGs specific to a given sequential comparison presented in each spreadsheet.

### Additional file 16: List of genes differentially expressed uniquely in one regeneration stage compared to all the others

Contain for each stage, separated by spreadsheet, the list of genes that are unique as well as their mouse annotation.

### Additional file 17: Tests of over-representation of functional categories within DEGs and clusters

-Enrichments for RNA-binding proteins (RBP), transcription factors (TF), nervous system genes (NS), and stem cell genes (SC) are tested for each DEGs lists for sequential comparisons (Spreadsheet 1), as well as the 12 clusters gene lists (Spreadsheet 2).

Spreadsheet 1: The total number of annotated transcripts for the Reference transcriptome, the number of DEGs per comparison, as well as the total number of TF, RBP, NS and SC gene searched are provided. The number and percentage of DEGs belonging to each category searched, as well as the p-valued are shown. Significant over-representations are highlighted in green.

Spreadsheet 2: The total number of transcripts / annotated transcripts per cluster are provided. The number and percentage of genes *per* cluster belonging to each category searched, as well as the p-valued are shown. Significant over-representations are highlighted in green.

### Additional file 18: Expression characteristics for a selection of known neurogenesis and stem cell genes for *Platynereis dumerilii*

Spreadsheet 1: For each gene associated to neurogenesis, the identifiers, median expression levels during the course of regeneration, as well as the sequential conditions in which the respective genes are up (blue) or down (red)-regulated are shown.

Spreadsheet 2: For each gene associated to stem cells, the identifiers, median expression levels during the course of regeneration, as well as the sequential conditions in which the respective genes are up (blue) or down (red)-regulated are shown.

### Additional file 19: GO term analysis of gene expression clusters

Treeplot representation of clustered enriched GO term (Biological process) analysis for 8 clusters (1, 2, 4, 7, 8, 9, 10 and 11). Circles area are proportional to the number of genes in each annotation, the colors represent the p-value of the over-representation.

### Additional file 20: Lists of DEGs belonging to each of the 12 clusters

Full list of gene identifiers for all DEGs belonging to any cluster, as well as their respective mouse annotation, are provided.

## Ethics approval and consent to participate

Not applicable

## Consent for publication

Not applicable

## Availability of data and materials

The datasets generated during the current study are available in the ENA repository. All persistent web links to datasets and accession numbers are provided in the Additional file 2.

## Authors’ contributions

EG and MV designed the study with the contribution of YC, LL, and SQS. LP performed most of data analysis with the help of YC. MB produced the ONT long read data. LBi, LBa and CD performed *in situ* hybridization experiments and imaging. EG, YC and LP wrote the manuscript with the contribution of LBi. EG, MV and LL obtained funding. All authors read and approved the final manuscript.

## Acknowledgments

This study is dedicated to the memory of our friend and mentor, the late Professor Michel Vervoort. We are indebted to all members of the Gazave & Vervoort lab for their support and suggestions on this study, especially Zoé Velasquillo for her help with biological material production, as well as Luis Carreira for his participation to initial ISH experiments. We are grateful to Dr. Jerome H.L. Hui and Wenyan Nong for their critical input in early stages of this study. We also thank the D. Arendt’s laboratory for sharing the *Otp* and *Brachyury* clones. We thank Corinne Blugeon for her input on the method section.

