## Supplementary figures and images for "Transcriptomic landscape of posterior regeneration in the annelid *Platynereis dumerilii*"

### Additional File 1

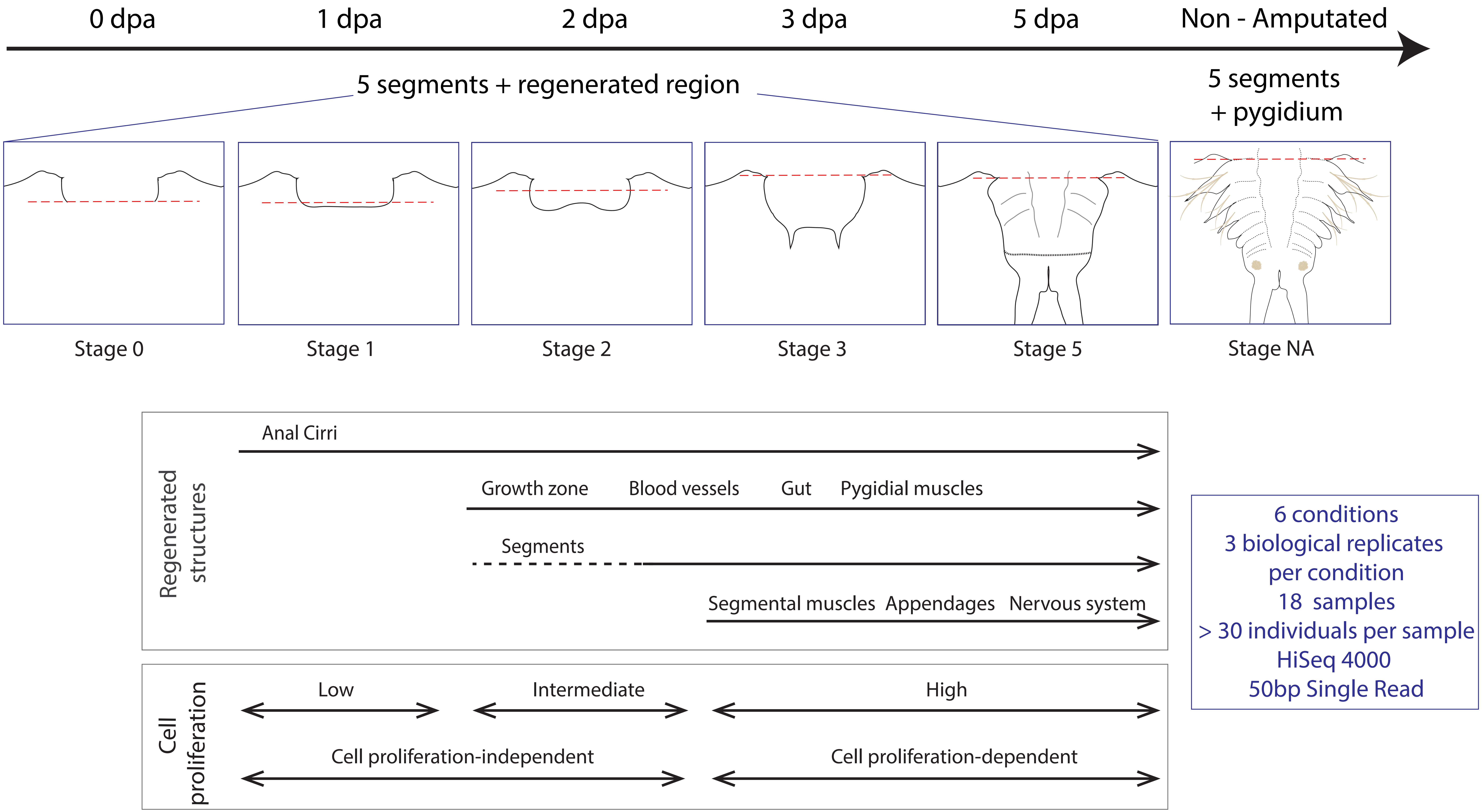

### Additional File 3

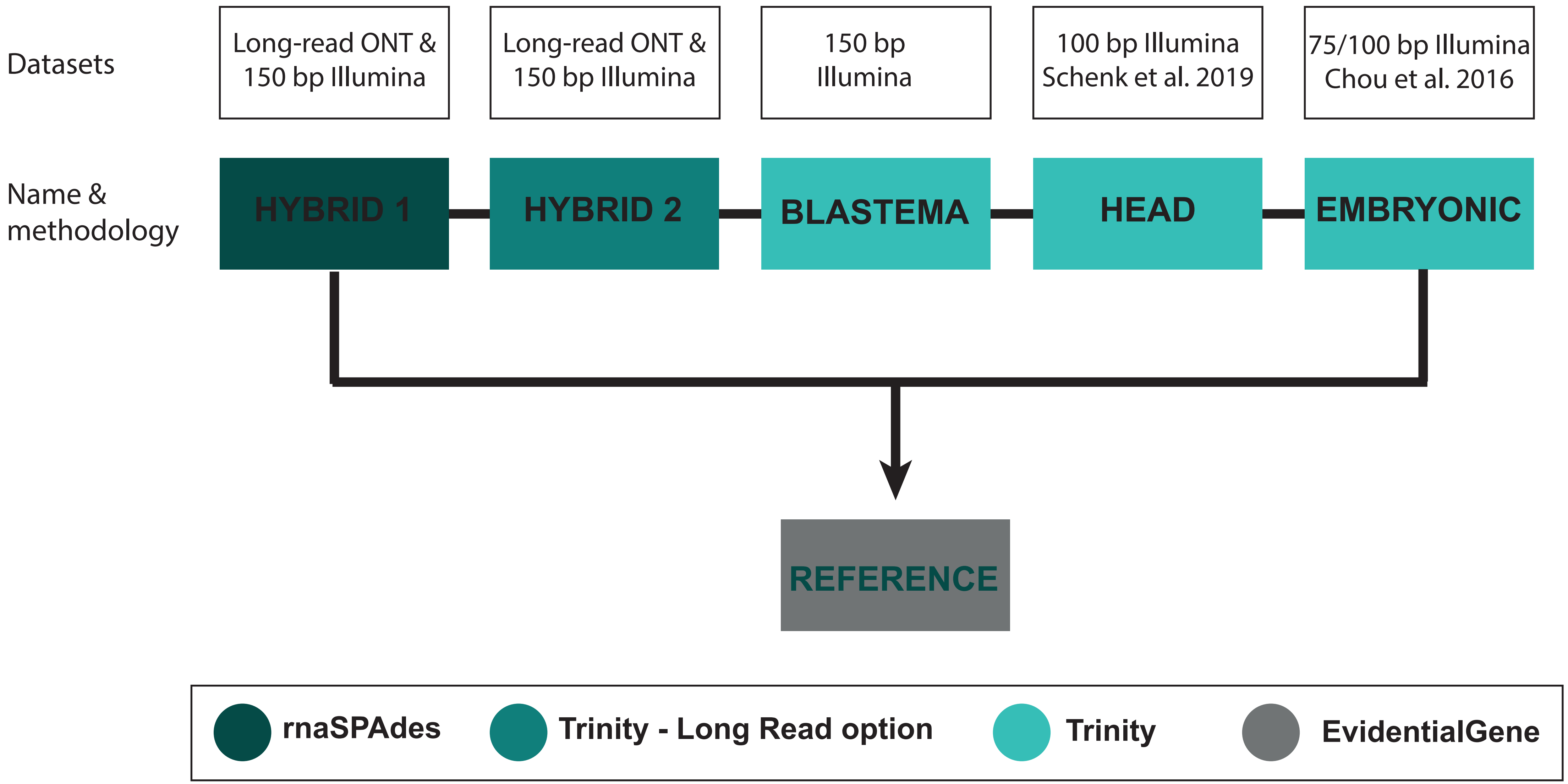

### Additional File 8

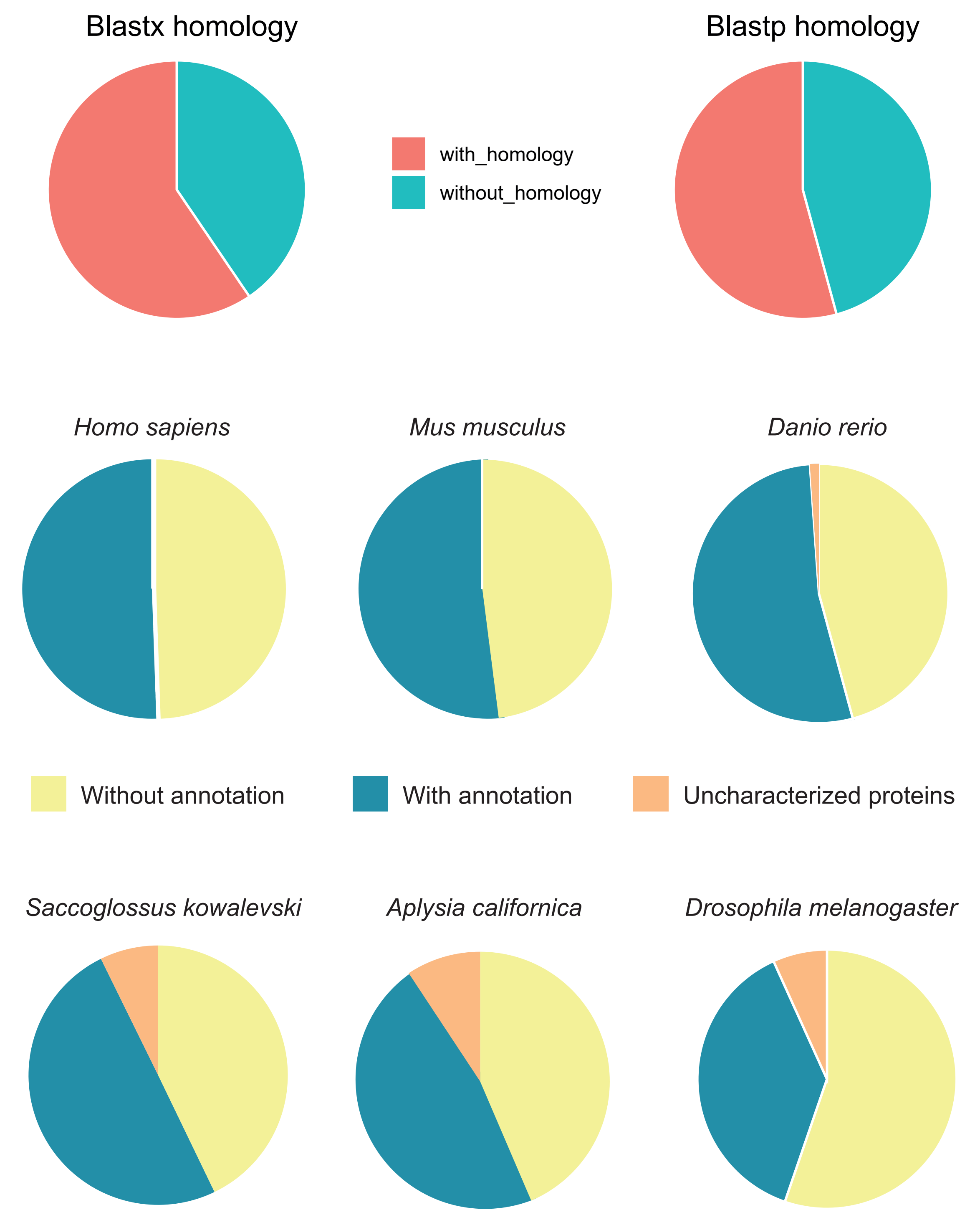

### Additional File 10

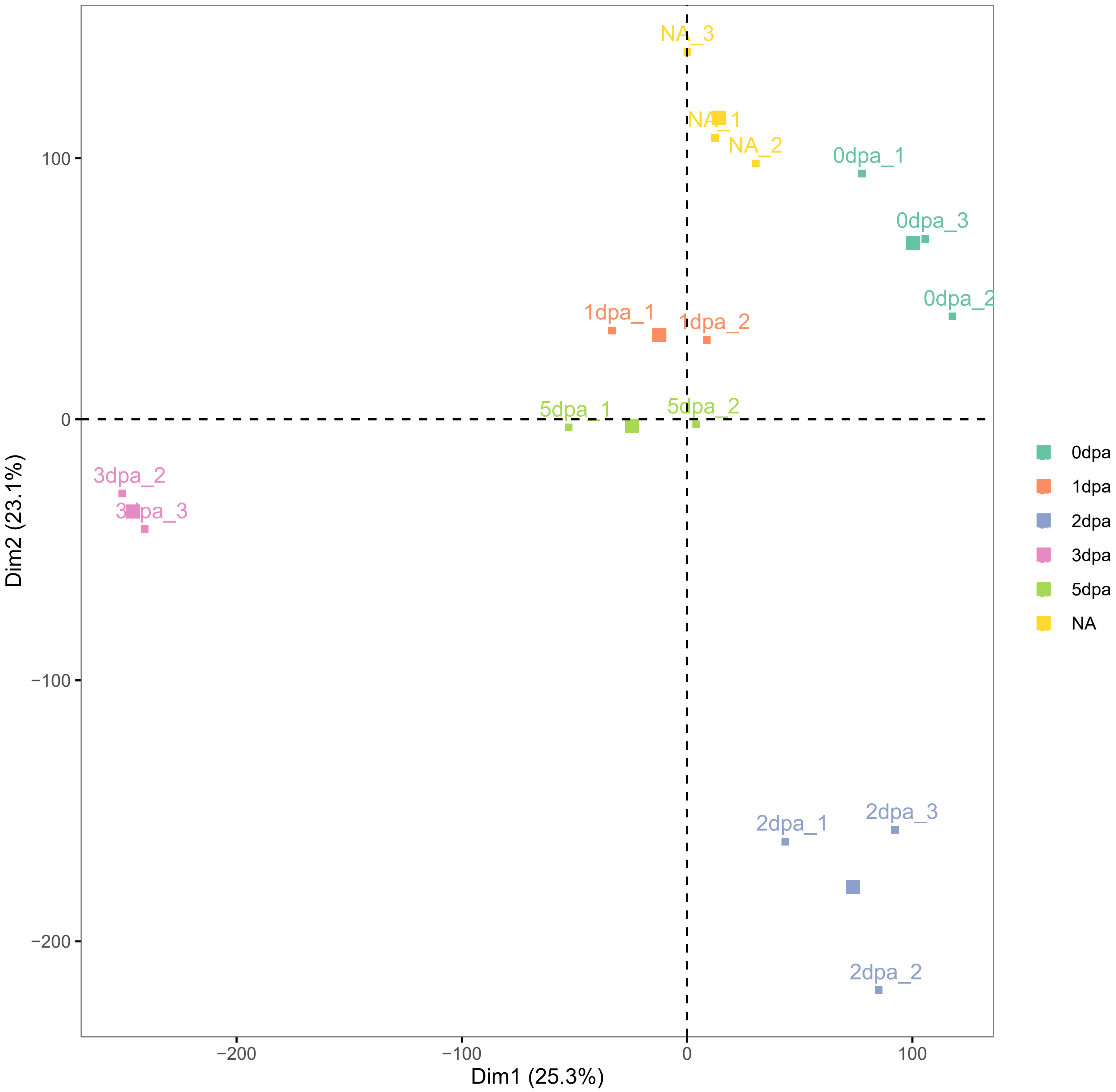

### Additional File 19

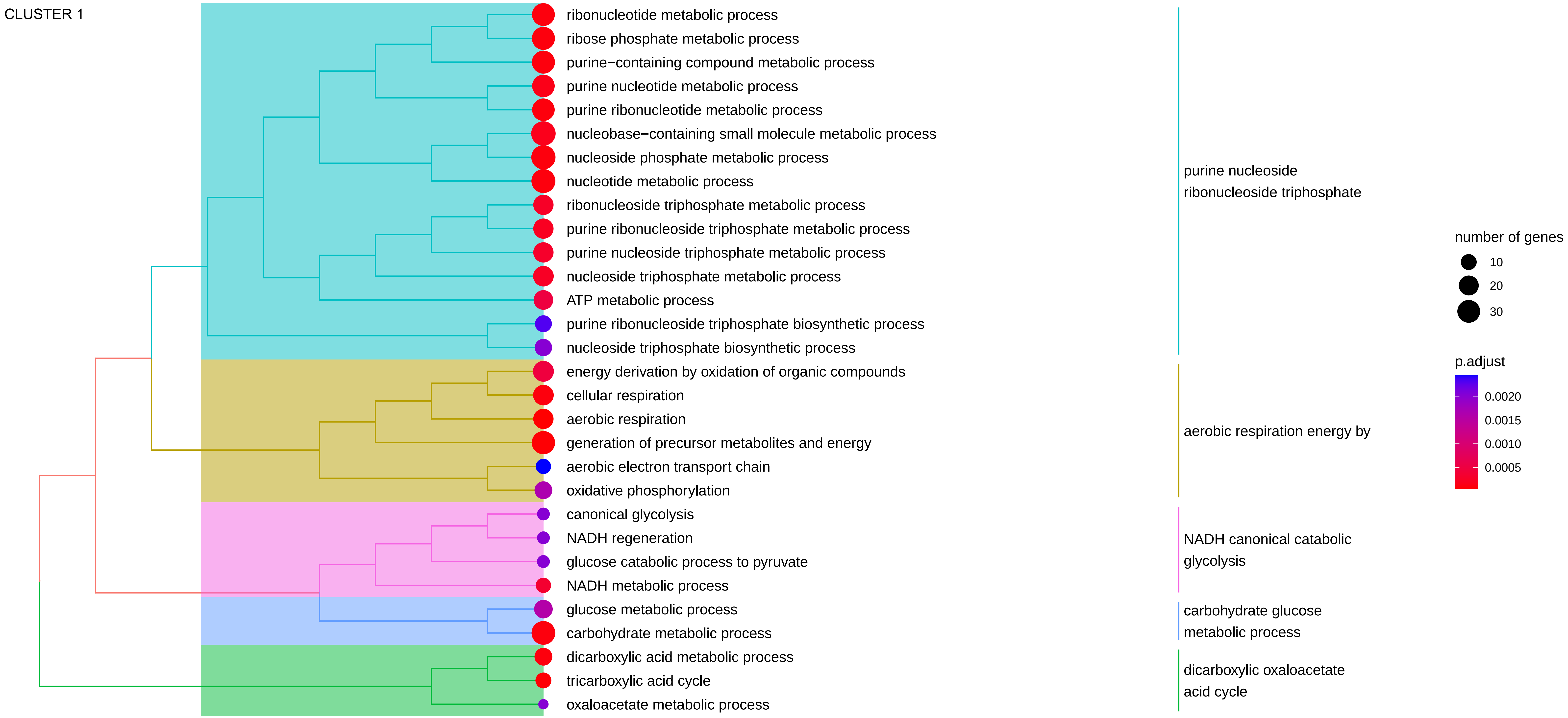

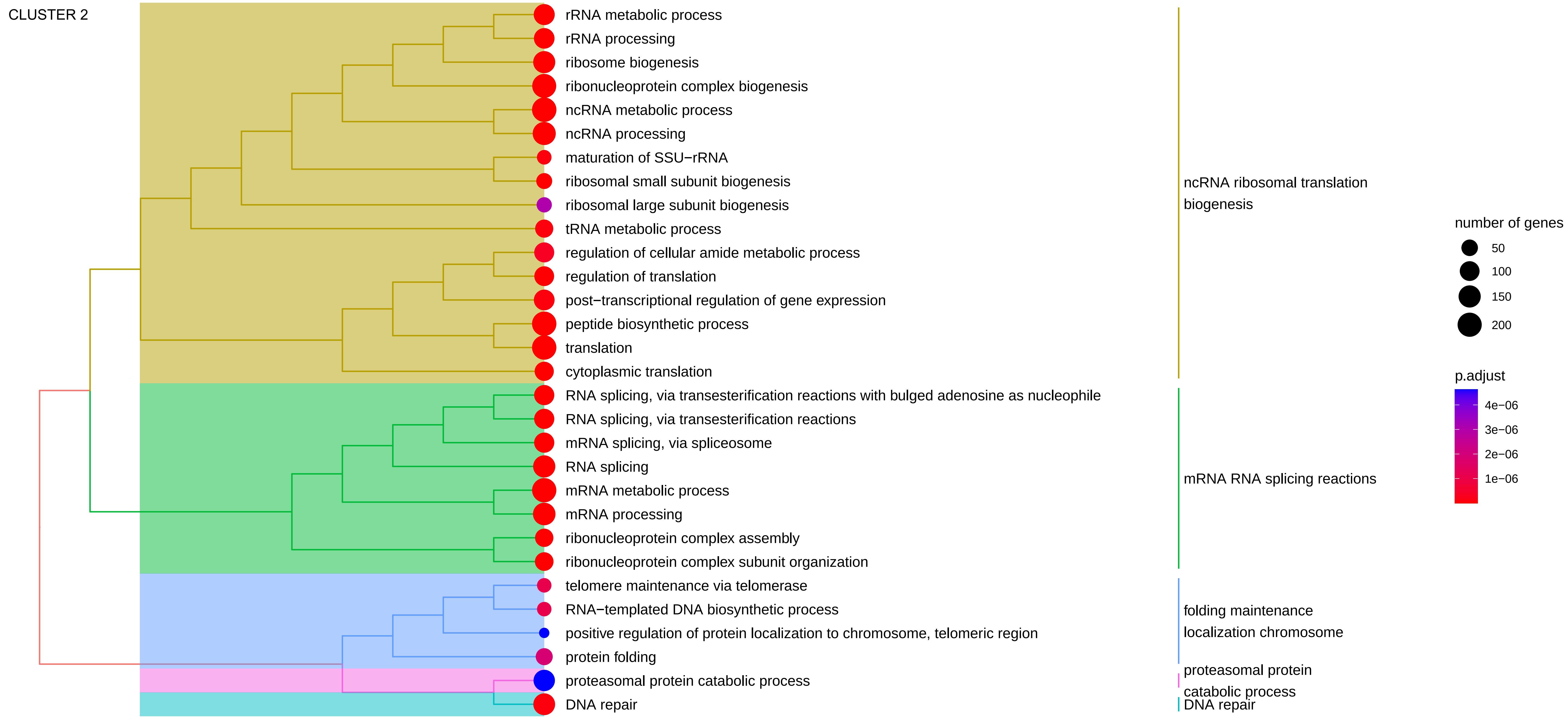

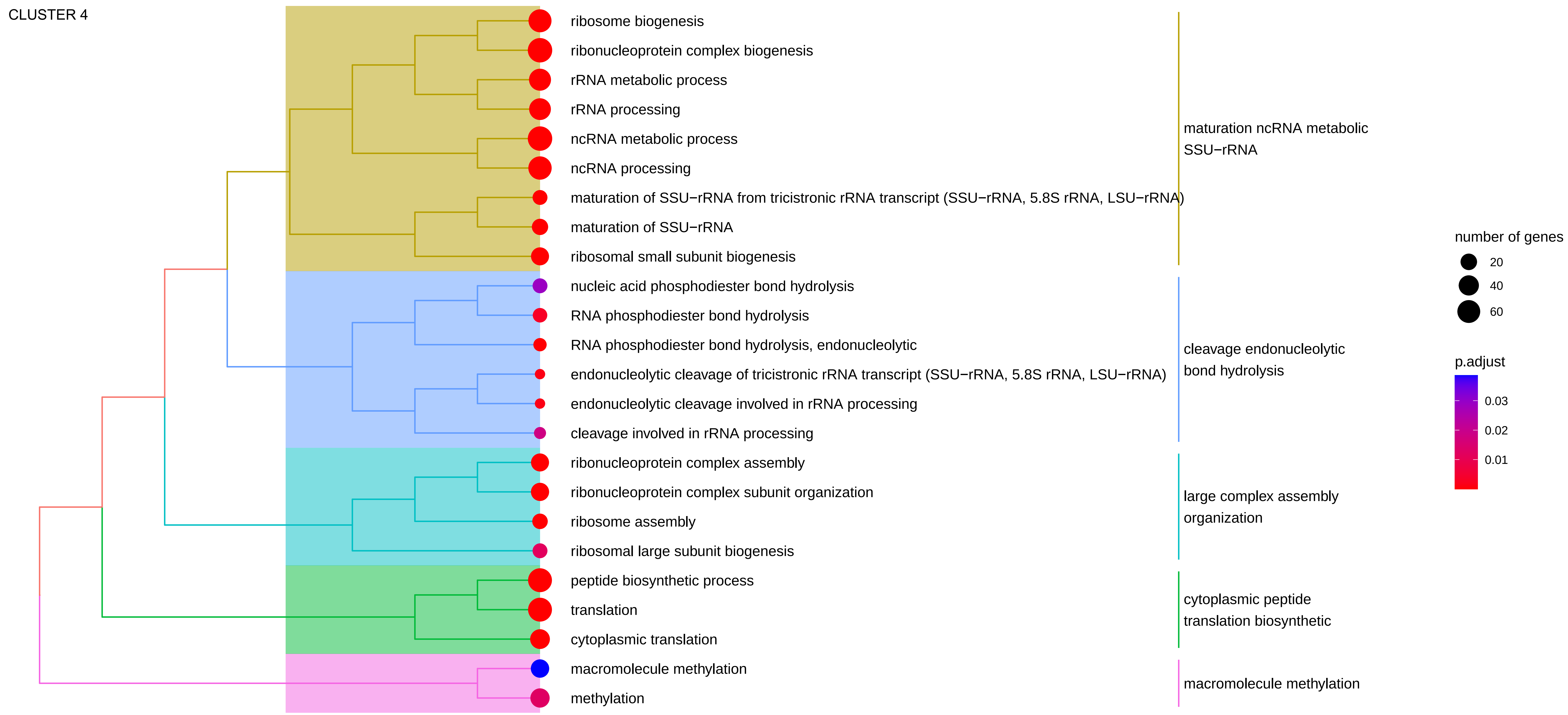

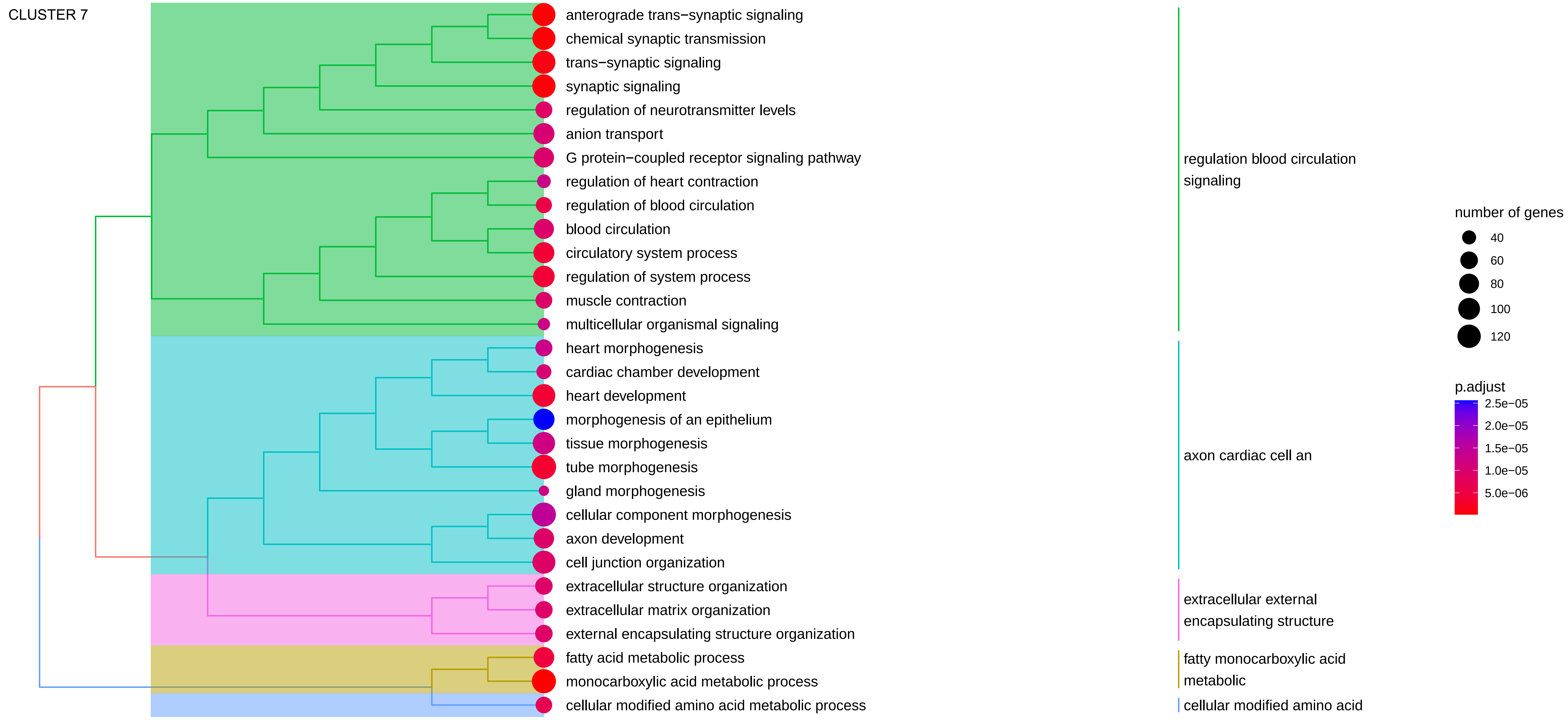

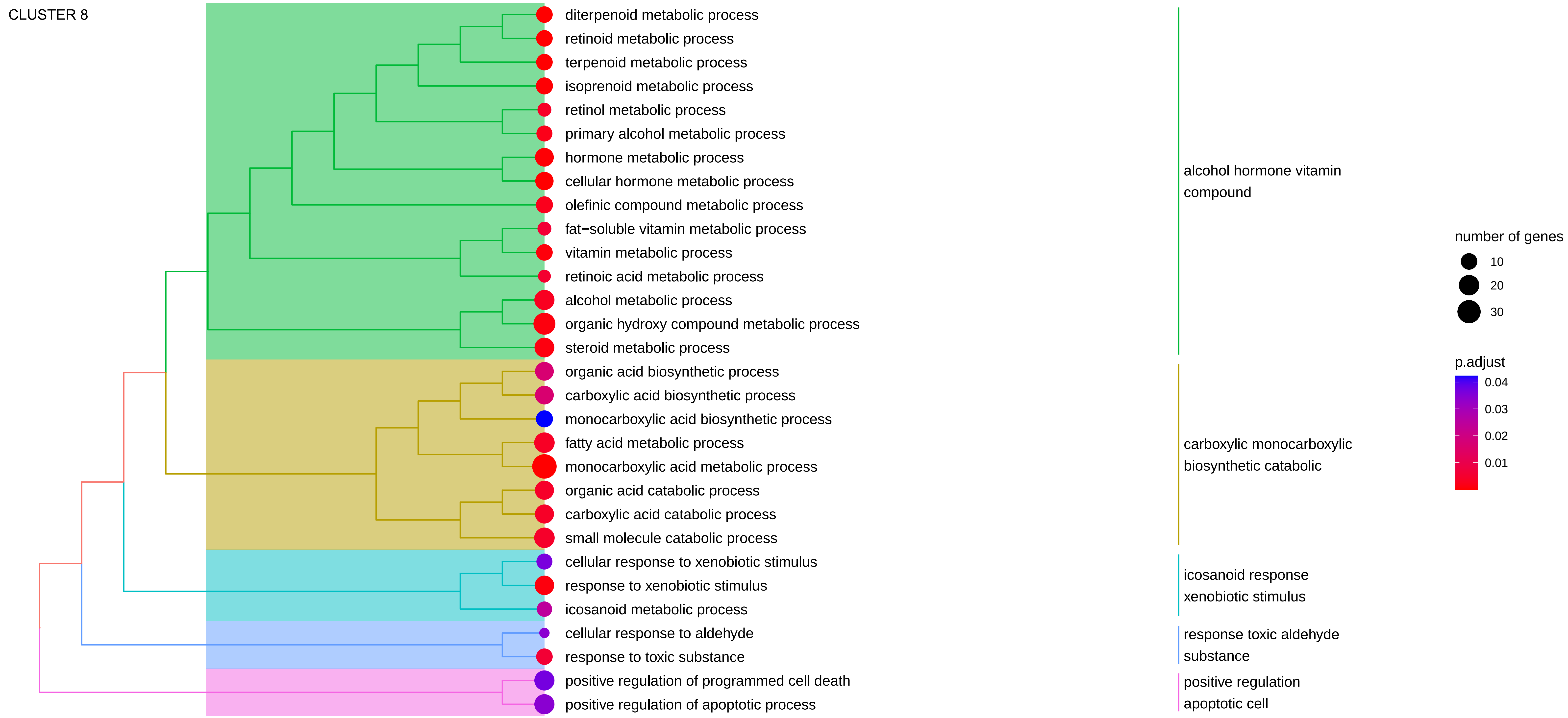

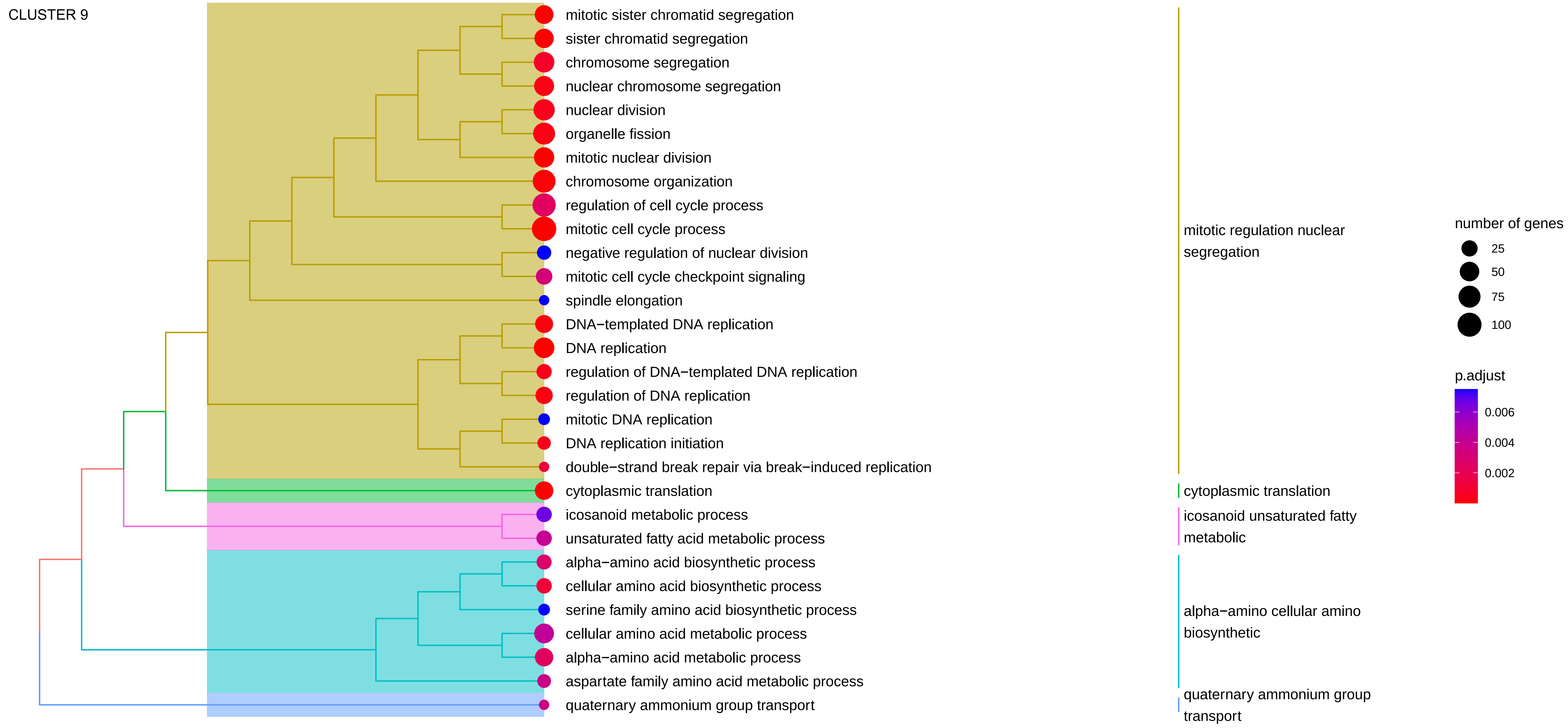

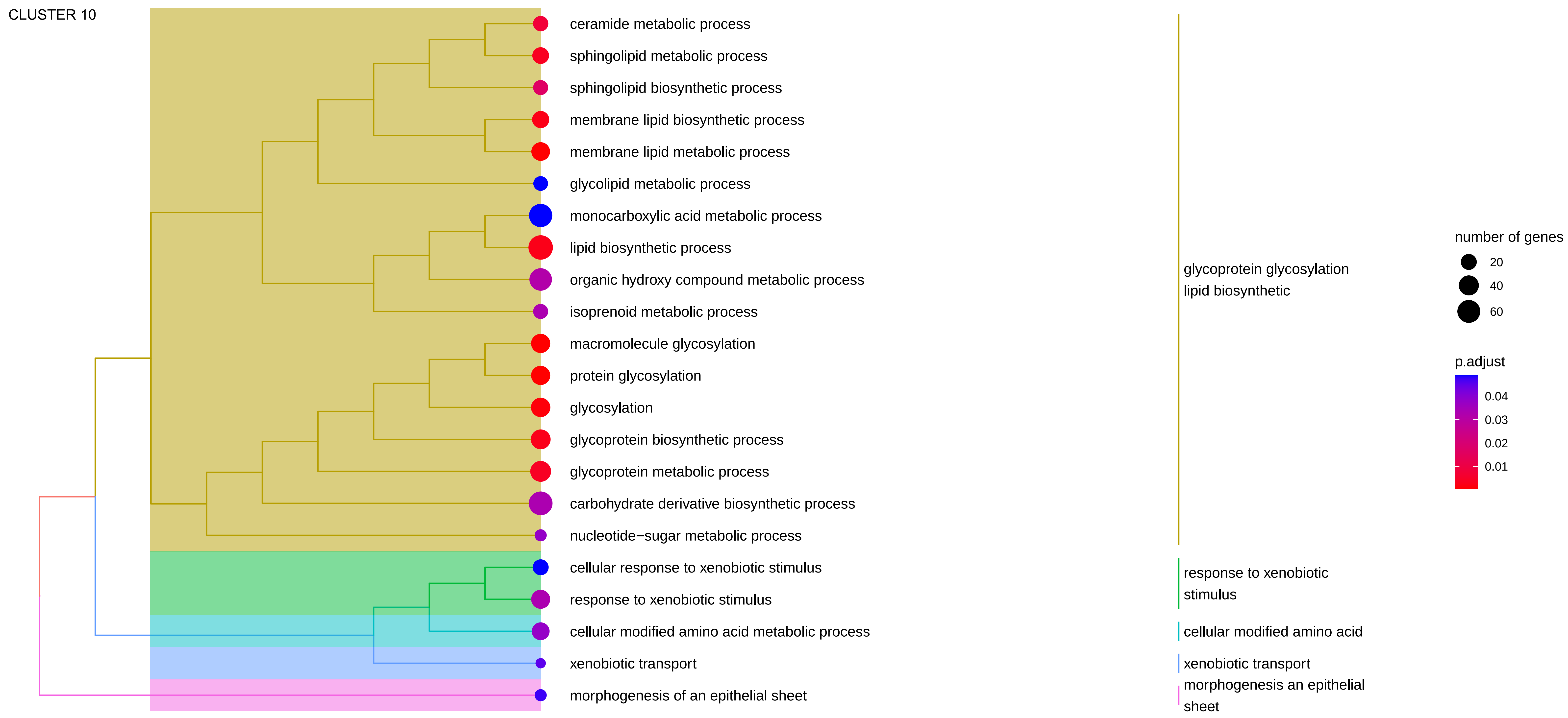

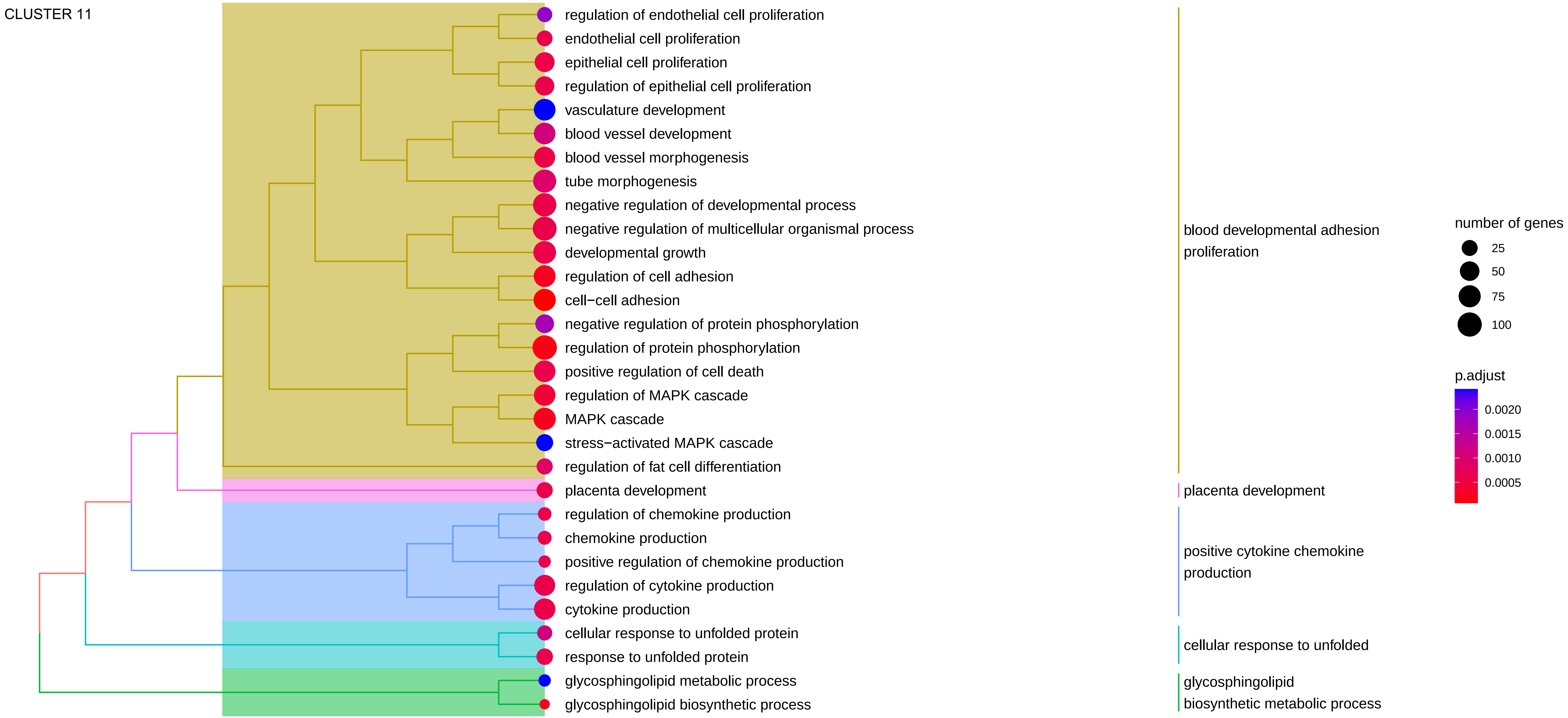
